# Single Cell Gene Expression Analysis Reveals Human Stem Cell-Derived Graft Composition in a Cell Therapy Model of Parkinson’s Disease

**DOI:** 10.1101/720870

**Authors:** Katarína Tiklová, Sara Nolbrant, Alessandro Fiorenzano, Åsa K. Björklund, Yogita Sharma, Andreas Heuer, Linda Gillberg, Deirdre B. Hoban, Tiago Cardoso, Andrew F. Adler, Marcella Birtele, Hilda Lundén-Miguel, Nikolaos Volakakis, Agnete Kirkeby, Thomas Perlmann, Malin Parmar

## Abstract

Since the pioneering studies using fetal cell transplants in Parkinson’s disease (PD), brain repair by cell replacement has remained a long-standing and realistic goal for the treatment of neurodegenerative disorders including PD. Authentic and functional midbrain dopamine (DA) neurons can now be generated from human pluripotent stem cells (hPSCs) via a floor plate intermediate^1,2^, and these cell preparations are both safe and functional when transplanted to animal models of PD^3^. However, although resulting grafts from fetal brain tissue and hPSCs contain large numbers of desired DA neurons, these therapeutic cells are a minor component of the grafts. Moreover, the cellular composition of the graft has remained difficult to assess due to limitations in histological methods that rely on pre-conceived notions concerning cell types. Here, we used single cell RNA sequencing (scRNA-seq) combined with comprehensive histological analyses to characterize intracerebral grafts from ventral midbrain (VM)-patterned human embryonic stem cells (hESCs) and VM fetal tissue after long-term survival and functional maturation in a pre-clinical rat model of PD. The analyses revealed that while both cell preparations gave rise to neurons and astrocytes, oligodendrocytes were only detected in grafts of fetal tissue. On the other hand, a cell type closely resembling a class of newly identified perivascular-like cells was identified as a unique component of hESC-derived grafts. The presence of these cells was confirmed in transplants from three different hESC lines, as well as from iPSCs. Thus, these experiments have addressed one of the major outstanding questions in the field of cell replacement in neurological disease by revealing graft composition and differences between hESC- and fetal cell-derived grafts, which can have important implications for clinical trials.

---

To compare the developmental potential of hESC-derived VM progenitors with fetal VM cells after transplantation in a rat model of PD, hESCs were subjected to VM-patterning by a well-established protocol intended for generation of clinical-grade cell preparations^4^ and human fetal tissue was dissociated from the VM of a 7.5 week old human embryo by the same protocol used for the fetal cell clinical transplantation trial, TRANSEURO^5^. Both types of cells were transplanted into the striatum of adult rats that had been unilaterally lesioned with 6-hydroxydopamine (6-OHDA) (Fig. 1a). Both fetal and hESC-derived cell preparations gave rise to grafts rich in DA neurons expressing tyrosine hydroxylase (TH) at six months following transplantation (Fig. 1b). In addition, paw use and rotational asymmetry induced by 6-OHDA lesioning was corrected in animals transplanted with hESC-derived VM progenitors, confirming a successful patterning of the cells prior to grafting and functional maturation after transplantation (Fig. 1c and d).

**Figure 1.**
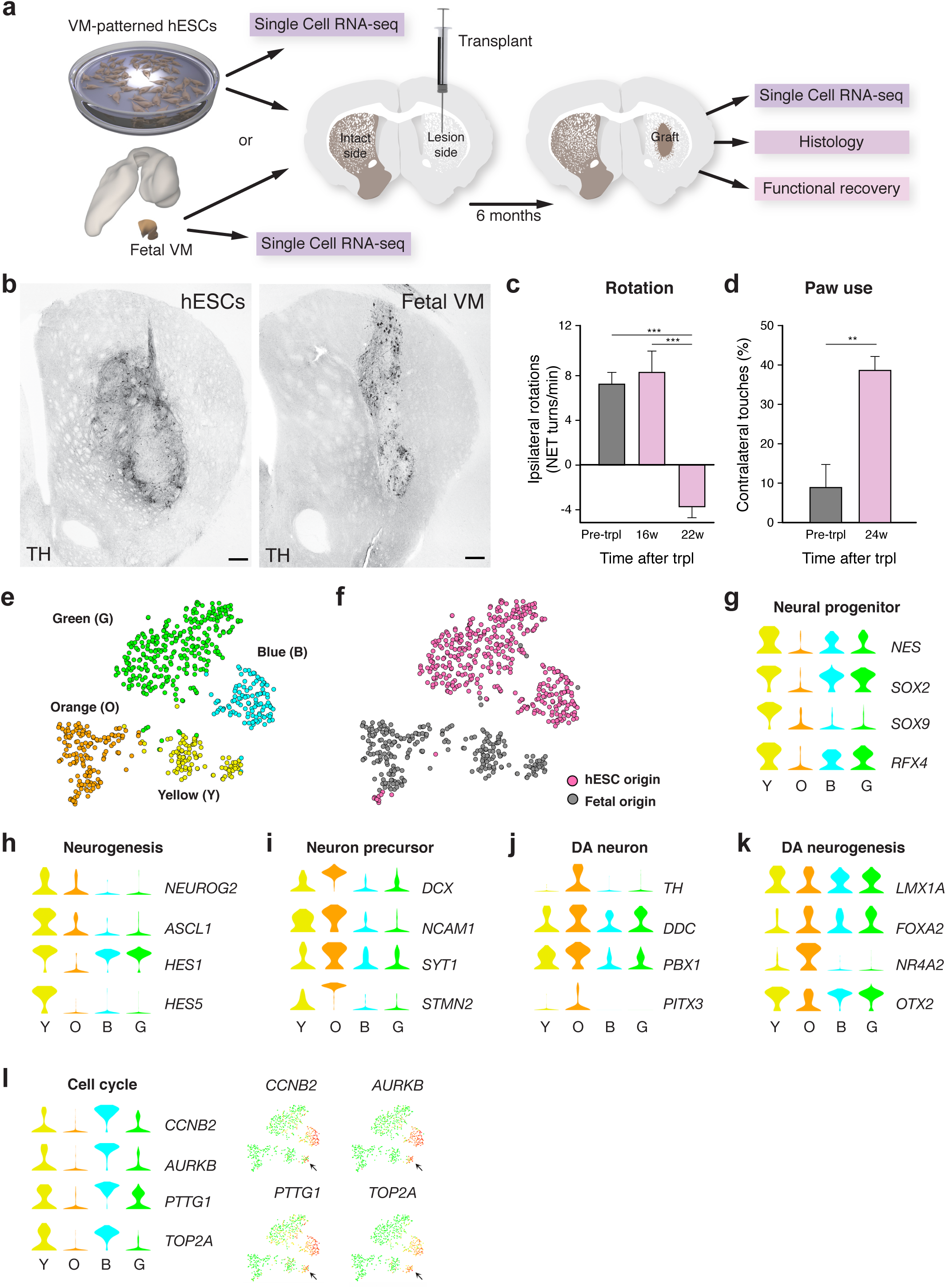
Histological validation and scRNA-seq analysis of progenitor cells before grafting in pre-clinical cell therapy model of PD. **a**, Schematic overview of experimental design. VM-patterned hESCs were grafted to seven rats and used as follows: scRNA-seq n=2; histology n=3; functional recovery n=7. Cells from one fetal VM were grafted to three rats and used as follows: scRNA-seq n=2; histology n=1. **b**, Immunohistochemistry of TH in hESC- and fetal VM-derived intrastriatal grafts 6 months post-transplantation. **c-d**, Functional recovery of the hESC-derived cells by amphetamine-induced rotation test and spontaneous paw use (Cylinder) test (n= 7 rats; mean ± SEM; p < 0.05 compared to post-lesion; two-tailed paired t-test).**e**, t-SNE showing clustering of analyzed cells before grafting. **f**, Same t-SNE as in e) but with origin of cells marked in pink (hESC) or grey (fetal) as indicated. **g-l**, Expression level per cluster for indicated genes. Genes represent markers for the cell types (neural progenitor, neuron precursor, DA neuron) or indicated processes (cell cycle, neurogenesis or DA neurogenesis; see text for details). Expression levels of indicated cell cycle genes are also shown in the t-SNE. Scale bar, 250 µM.

Cells were sampled for scRNA-seq both before grafting and 6 months after transplantation as illustrated in Figure 1a. To separate intact cells from cell fragments and debris, fetal and hESC-derived progenitors were isolated by fluorescence activated cell sorting (FACS) based on cell size (forward and side scatter) before grafting (Extended Data Fig. 1a). t-distributed neighbor embedding (t-SNE) and graph-based clustering of the scRNAseq data from fetal and hESC-derived cells before grafting resulted in four major clusters (green, blue, orange and yellow in Fig. 1e). The two top clusters (green, blue) consisted almost exclusively of VM-patterned hESCs while the two bottom clusters (orange, yellow) consisted mostly of fetal cells (Fig. 1f), suggesting that while both fetal and hESC-derived cells generate DA neurons with similar subtype specificities and functional properties after grafting^6,7^, the two different sources of cells were transcriptionally distinct at the time of transplantation.

Samseq was used to identify genes enriched in each cluster (Extended Data Table 1; Extended Data Fig. 2a, b). Neural progenitor/radial glial cell markers including *NES, SOX2, SOX9* and *RFX4* were prominently expressed in the yellow fetal cell-dominated cluster as well as in the blue and green hESC-dominated clusters consistent with the expression of these genes also under DA neurogenesis in mouse and human^8,9^ (Fig. 1g; Extended Data Fig. 2c). Regulators of neurogenesis (*NEUROG2, ASCL1, HES1* and *HES5*) were prominently expressed in the yellow fetal cell-dominated cluster while the blue and green hESC-dominated clusters mainly expressed *HES1*^8,9^ (Fig. 1h; Extended Data Fig. 2d). Importantly, transcription factors such as *LMX1A, FOXA2* and *OTX2* that are critical in ventral midbrain development and DA neurogenesis, as well as genes that are predictive of successful graft outcome (*EN1* and *DLK1*)^10,11^, were expressed by both fetal and hESC-derived cells (Fig. 1k, Extended Data Fig. 2e). Cells in the fetal cell-dominated orange cluster (Fig. 1e) expressed genes that are normally active in early developing neurons including *DCX, NCAM1, SYT1* and *STMN2*, as well as genes that are normally expressed in maturing DA neurons including *NR4A2, TH, DDC, PITX3* and *PBX1* (Fig. 1i, j, k; Extended Data Fig. 2e, f, g). Of note, cells expressing markers for more mature DA neurons could be distinguished in one part of the orange cluster, thus indicating neuronal diversity among more mature fetal cells (Extended Data Fig. 2g).

Expression of cell cycle genes and scores that assigned cells to different cell cycle phases indicated the presence of cycling cells within the blue (hESCs) and yellow (fetal) clusters (Fig. 1l; Extended Data Fig. 3a). From the list of enriched genes it is evident that the difference between the green and blue clusters correspond to cell cycle genes being expressed only in the blue cluster (Extended Data Table 1). Markers of pluripotency (*POU5F1, NANOG*), mesoderm (*T*) or endoderm (*SOX17*) development were not expressed in any of the cell clusters (Extended Data Fig. 2h, i).

Six months after transplantation, when grafts had reached functional maturity, the transplants were dissected from the striatum, dissociated into a single cell suspension and subjected to FACS (see Fig 1a, Extended Data Fig. 1b). The transplanted hESCs contained a GFP-encoding transgene expressed in virtually all transplanted cells (Extended Data Fig. 1e, g). Fetal cells did not carry a GFP-encoding transgene and were therefore isolated by FACS based on cell size (forward and side scatter) to separate single cells from debris (Fig. 1a, Extended Data Fig. 1a). Notably, since fetal cells did not express GFP, the vast majority of single cells isolated from fetal grafts were contaminating rat cells and scRNA-seq data from fetal transplants was therefore limited to only 63 fetal cells. In contrast, several hundred cells in each condition could be analyzed from hESC-derived grafted cells as well as from both fetal- and hESC-derived cells before grafting (Extended Data Fig. 1d).

Using graph-based clustering, fetal and hESC-derived cells sampled from rat striatum six months after grafting segregated into four main clusters (Fig. 2a). We used Samseq^12^ for the identification of genes enriched in each cluster (see Methods; Extended Data Table 2; Extended Data Fig. 4b). From inspection of the most highly enriched genes in each cluster it was evident that three of the clusters contained cells expressing genes characteristic of astrocytes (*AQP4* and *GFAP)*, oligodendrocytes (*OLIG1* and *OLIG2*) and neurons (*GAP43* and *RBFOX3)*, respectively (Fig. 2b-d, Extended Data Fig. 4c-e). Of note, *PMP2*, one of the most significantly enriched genes in the oligodendrocyte cluster is a marker for peripheral Schwann cells in mouse but was recently shown to be prominently expressed in human oligodendrocytes^13^. In addition, dopaminergic markers including *TH, SLC18A2* and *DDC* were expressed in cells in the neuron cluster indicating, as expected, a high proportion of DA neurons (Fig. 2e, Extended Data Fig. 4f). Further analyses of publicly available data sets reporting both bulk and scRNA sequencing provided additional strong support for the assignment into astrocytes, oligodendrocytes and neurons^13-16^ (Extended Data Fig. 5a). Of note, despite transplantation of the cells at a highly proliferative stage, only a very small number of cells (1.3%) showed cell cycle scores indicative of some cycling cells after six months *in vivo*, and these cells all belonged to the oligodendrocyte and astrocyte clusters (Extended Data Fig. 3b).

**Figure 2.**
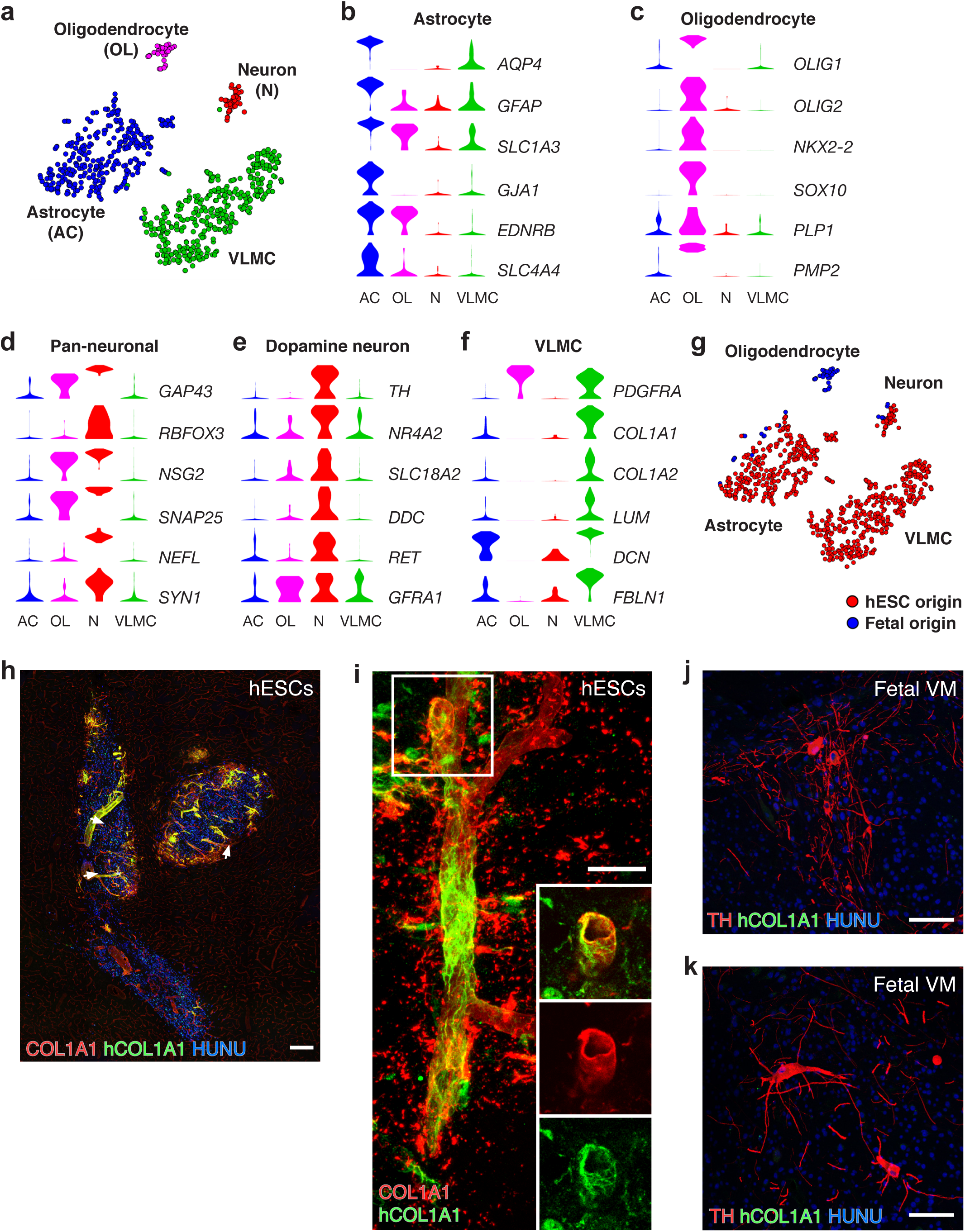
scRNA-seq analysis and histological validation of grafted cells into the striatum. **a**, t-SNE showing clustering of analyzed intrastriatal grafted cells. Cell type assignments are indicated. **b-d**, Expression level per cluster for indicated genes. All indicated genes are significantly enriched and established markers for astrocytes, oligodendrocytes and pan-neuronal cells, respectively. **e**, Expression level per cluster for selected dopaminergic markers. **f**, Expression level per cluster for genes that are significantly enriched in the VLMC cluster and established markers for barrier forming fibroblasts including VLMCs. **g**, t-SNE of grafted cells as shown in Fig. 2a. Cells are marked according to their origin from either hESC-derived (red) or fetal-derived (blue) transplants. **h**, Staining using antibodies recognizing both rat and human COL1A1 or only hCOL1A1 as indicated in a hESC-derived grafts from same experiment as used for RNAseq. **i**, Representative immunofluorescence micrograph of hCOL1A1-positive cells intermingled with host-derived COL1A1-positive cells in close association with blood vessels. Human nuclei were counterstained with HUNU. **j-k**, Representative micrographs of TH/hCOL1A1 immunostaining in fetal derived grafts from the same experiment as used for RNAseq (j) and from a different experiment using fetal tissue from another fetal VM of same gestational age. Scale bars, 200 µM (h), 20 µM (i) and 100 µM (j,k)

It is notable that the overall proportion of neurons relative to other cell types was remarkably low (<10%) in the scRNA-seq analysis from both fetal- and hESC-derived grafts (Fig. 2a). This finding significantly diverged from the high neuronal content observed in previous histological analyses of rats grafted with VM-patterned hESCs^2,10,17-19^. We therefore further investigated the neuronal content in brain sections of animals grafted at the same time with the same preparation of hESCs/fetal cells that were used for sequencing. In contrast to the scRNA-seq data, a high neuronal content and extensive innervation that matches previous results^2,7,10^ was observed also in these grafts (Extended Data Fig. 5b and c). The neuronal content of the hESC-derived grafts was further confirmed by comparing the number of NEUN^+^ cells from this experiment as well as a representative separate experiment where a different preparation of cells was transplanted in a separate grafting session but according to the same procedure (Extended Data Fig. 5d, e). The apparent underrepresentation of neurons among recovered cells in the scRNA-seq data was not surprising considering difficulties to achieve complete mechanical dissociation of brain tissue into single cell suspensions of mature neurons containing extensive projections^14,20^. Our data therefore underscores the importance of validating both qualitative and quantitative results from scRNA-seq data sets by alternative methods.

The scRNA-seq data also identified a fourth cluster of cells which could not, based on the top enriched genes, be easily assigned to a known neural cell type (Extended Data Table 2; Extended Data Fig. 4b). Gene enrichment analysis using the top distinguishing genes of this cluster identified highly significant biological functions associated with vasculature and connective tissue, including genes encoding collagens and other extracellular components (Fig. 2f; Extended Data Fig. 6a and Extended Data Table 3). We then probed a large scRNA-seq data set from the mouse brain interrogating more than 400,000 cells organized into 250 annotated clusters representing diverse cell types such as central and peripheral neurons, astrocytes, oligodendrocytes, Schwann cells, immune cells, and brain vascular cell types^14^ (Mouse Brain Atlas, http://mousebrain.org/). Interestingly, when this data set was searched for cell types co-expressing enriched genes of the fourth cluster, “enteric glia” and “vascular leptomeningeal cells” (VLMCs) were identified as cell type categories with highly related gene expression profiles (Extended Data Fig. 6b, c). Enteric mesothelial fibroblasts, the closest matching cell type among “enteric glia”, and VLMCs belong to a family of highly related organ barrier-forming fibroblast populations^14^. Recent scRNA-seq analyses have identified related cells associated with the mouse brain vasculature^15,21^, characterized by the co-expression of *Pdgfra, Col1a1, Col1a2, Lum* and *Dcn*^14,15,21^. Interestingly, the human homologues of these genes were all expressed in the unknown cluster (Fig. 2f, Extended Data Fig. 4g). Therefore, we refer to these cells as VLMCs as this cluster consists of cells with striking transcriptional resemblance to previously described populations of barrier-forming fibroblasts, including VLMCs of the mouse central nervous system^14,15,21^.

The analysis of graft composition uniquely allowed us to discern similarities and potential differences between the cellular composition of fetal and hESC-derived grafts after transplantation. Strikingly, tracing the origin of grafted cells visualized by t-SNE suggested prominent differences in generated cell types (Fig. 2g). Accordingly, two of the clusters (astrocyte and neuron clusters) contained cells both from grafted VM-patterned hESCs and from fetal cells. In contrast, the oligodendrocyte cluster contained only fetal-derived cells while the VLMC cluster contained only hESC-derived cells (Fig. 2g). The scRNA-seq analysis of cells after grafting thus suggested that VM-patterned hESCs and fetal VM cells have distinct developmental potential after grafting. Based on the unexpected identification of VLMCs in the hESC grafts, we examined highly expressed genes found in the VLMC cluster (*COL3A1, FBLN1, S100A11* and *IFITM2)*, and additional genes encoding components of the extracellular space (e.g. *EMP2* and *MMP2*), also in the cells prior to grafting (Extended Data Fig. 6d). We found that several of these VLMC-associated genes were expressed together with DA progenitor markers in the same hESC-derived cells suggesting that a common hESC-derived precursor has potential for generation of neurons, astrocytes and VLMCs.

To validate the key observations from the scRNA-seq analysis, and to investigate in more detail the properties of graft-derived VLMCs, we first confirmed the presence of VLMCs in brain sections of animals grafted in parallel with the same batches of cells that were used for scRNA-seq. Using antibodies recognizing both rodent and human COL1A1 (uniquely expressed in the VLMC cluster among the transplanted cells), and an antibody that is specific for the human protein (hCOL1A1) we confirmed the presence of VLMCs in the hESC-derived grafts (Fig. 2h, i). These COL1A1^+^ cells in the hESC-derived grafts were commonly located in close association with blood vessels penetrating into grafts (arrows in Fig. 2h), which is consistent with previous studies describing VLMCs associated with the brain vasculature^14,15,21^. The analysis indicated that the hESC-derived VLMCs were intermingled with rat cells of similar appearance and location within the transplants (Fig 2h, i), suggesting that cells originating from both host and hESCs contributed to the graft vasculature as has been reported for rodent-to-rodent neural transplants^22,23^. Co-labeling of TH and hCOL1A1 in transplants from the same fetal cells grafted at the same time as the cells used for sequencing showed that very few if any hCOL1A1-expressing cells were present (Fig. 2j), and co-labeling of hCOL1A1/COL1A showed that the cells associated with the vasculature in these grafts were exclusively host derived (Extended Data Figure 7a). The lack of VLMCs in fetal VM grafts was confirmed in an additional transplant experiment using tissue from another 7.5 week old embryo (Fig 2k and Extended Data Fig. 7b), as well as transplants derived from a younger embryo (5.5 week old, Extended Data Fig. 7c) and in *in vitro* cultures from two additional embryos (Extended Data Fig. 7d, e). Also in agreement with the transcriptional analysis, graft derived oligodendrocyte progenitors expressing OLIG2 and PDGFRa were detected at the edge of fetal VM grafts (Extended Data Fig. 7f-h) but not hESC-derived grafts (Extended Data Fig. 7i). Thus, the histological analysis clearly confirmed the findings from scRNA-seq showing different potential to generate VLMCs and oligodendrocytes depending on the cellular origin of the transplanted cells.

To assess if the identification of VLMCs in the hESC-grafts apply also to other cell lines and differentiation protocols, we next assessed the presence of VLMCs in grafts from previous transplantation experiments using VM-patterned progenitors derived from three different hESC lines: RC17 (GMP protocol showing minimal variability^4^), H9 (research grade protocol) and HS980^4,6,7,10^ (GMP protocol) (Fig. 3a-c), and an iPSC line (GMP protocol) where the cells were sorted for DA progenitor marker prior to grafting (Fig. 3d)^24^. Importantly, graft derived hCOL1A1^+^ cells were present in all analyzed VM-patterned hPSC-derived grafts independent of cell line or differentiation protocol (Fig. 3a-d). Clinical delivery will be of cells that have been cryopreserved prior to transplantation, and VLMCs were shown to be present to a similar extent in grafts from fresh and cryopreserved cells (Fig. 3e, f).

**Figure 3.**
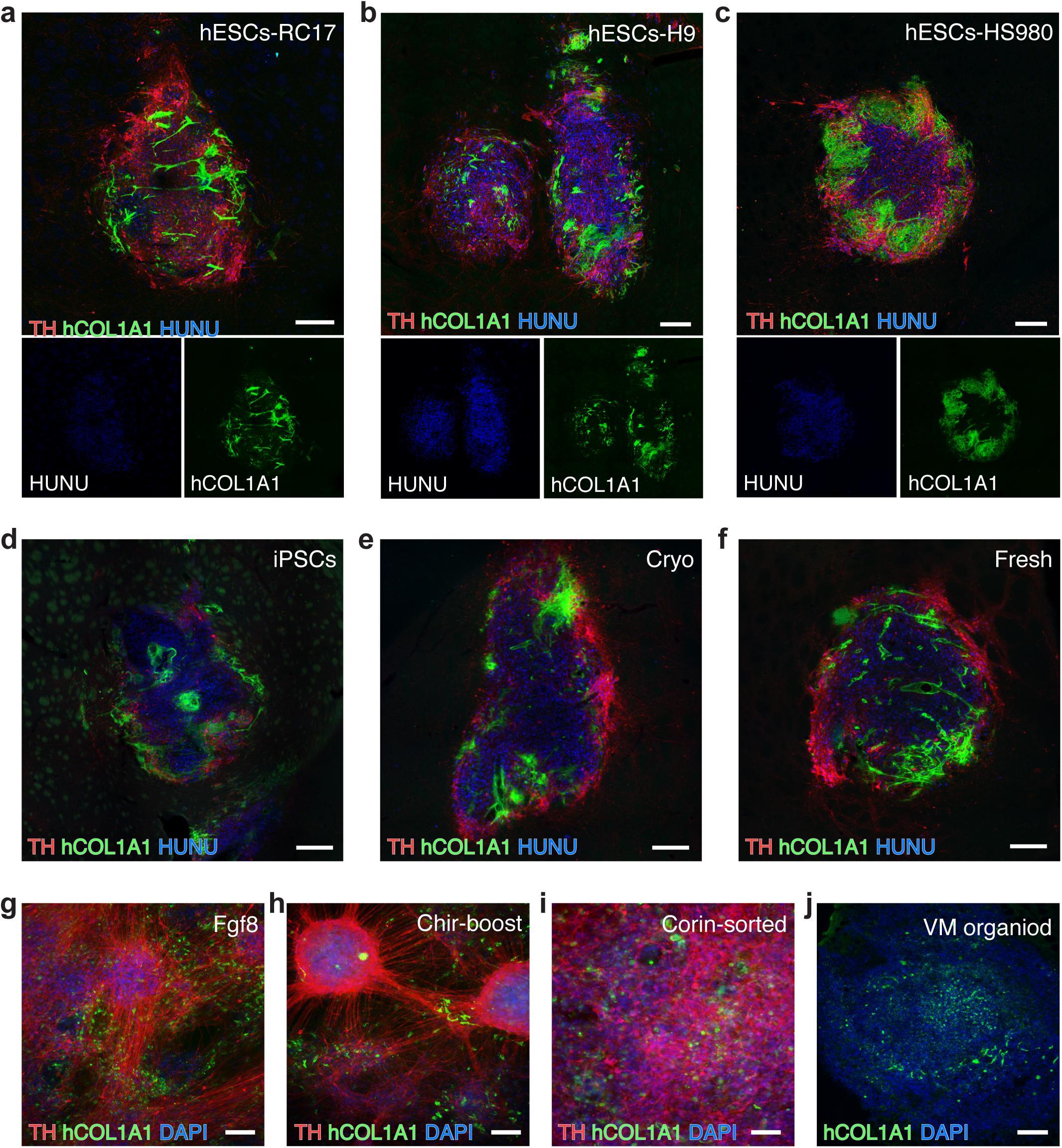
Presence of VLMCs in different cell lines. **a-c**, Representative immunofluorescence micrograph showing staining of hCOL1A1 in TH-rich hESC-derived grafts from three additional grafting experiments where the transplant was generated from VM patterned RC17 (a), H9 (b,) ^24^ and HS980 hESC lines (c) ^4^. **d**, TH/ hCOL1A1 immunofluorescence staining of a VM patterned iPSC-transplant derived sorted by IAP expression prior to transplantation^24^. **e-f**, TH**/** hCOL1A1 immunofluorescence staining of VM-patterned hESC-derived grafts generated from cryopreserved cells (e) or fresh cells (f). **g-i**, Representative micrographs of TH/hCOL1A1 double immunostaining in terminally differentiated hESC *in vitro* cultures derived by three different clinically relevant VM-patterning differentiation protocols: the protocol used in this study (g), a protocol developed in the Studer lab that uses CHIR boost instead of FGF8 for proper caudalization (described in https://patents.justia.com/patent/20180094242) (h), and a protocol developed by the Takahashi lab where the cells are sorted based on Corin prior to grafting (insert ref to Doi and Kukuchi) (i)). Nuclei were counterstained with DAPI in all three cultures **j**, hCOL1A immunostaining in a self-organized midbrain patterned organoid. Scale bars, 200 µM (a-f) and 100 µM (g-j).

To investigate if the derivation of VLMCs was a characteristic uniquely associated to the specific differentiation protocols used in this study, we assessed hESCs after terminal differentiation *in vitro* using two additional recently published protocols for generating VM progenitors from hPSCs^11,25^ see also https://patents.justia.com/patent/20180094242) as well as in three-dimensional self-organized organoids. These protocols all gave rise to cultures that are rich in TH^+^ neurons and also cells co-expressing VLMC markers COL1A1 (Fig. 3g-j) and PDGFRA (Extended Data Figure 7j-m).

The scRNA-seq analysis of grafted cells show that hESC-derived transplanted cells give rise to neurons, astrocytes and VLMCs, but not to oligodendrocytes. In these experiments detection of GFP (from the *SYN-GFP* transgene) was used to isolate grafted hESC-derived cells by FACS. To validate and extend the findings, hESCs patterned by the same protocol as initially used (Fig. 1 and 2) were grafted to the midbrain of nude rats and analyzed by scRNA-seq after 9 months of *in vivo* maturation (Fig. 4a-d). Importantly, to ensure that the GFP reporter and method of cell isolation did not influence/bias the results, cells were in the new experiment sequenced either directly after dissociation or after FACS isolation (outlined in Fig. 4a). In addition, the 10X Genomics Platform was used to allow for higher throughput. After QC and filtering to exclude rat cells (see Extended Data Fig. 8a-c), a total of 7875 cells were retained for analysis. The resulting UMAP embedding and graph-based clustering showed that, as with the hESC-derived cells grafted to the striatum (Fig. 2), VM-patterned hESCs transplanted to the midbrain gave rise to three main clusters which, based on marker expression, were clearly classified as astrocytes, neurons and VLMCs (Fig. 4e-i; Extended Data Table 4). A small number of cells expressing astrocyte markers expressed cycling genes and clustered separately (Fig. 4e). Similar results were derived regardless of whether cells had been isolated and sequenced directly or isolated by FACS before sequencing (Fig. 4j).

**Figure 4.**
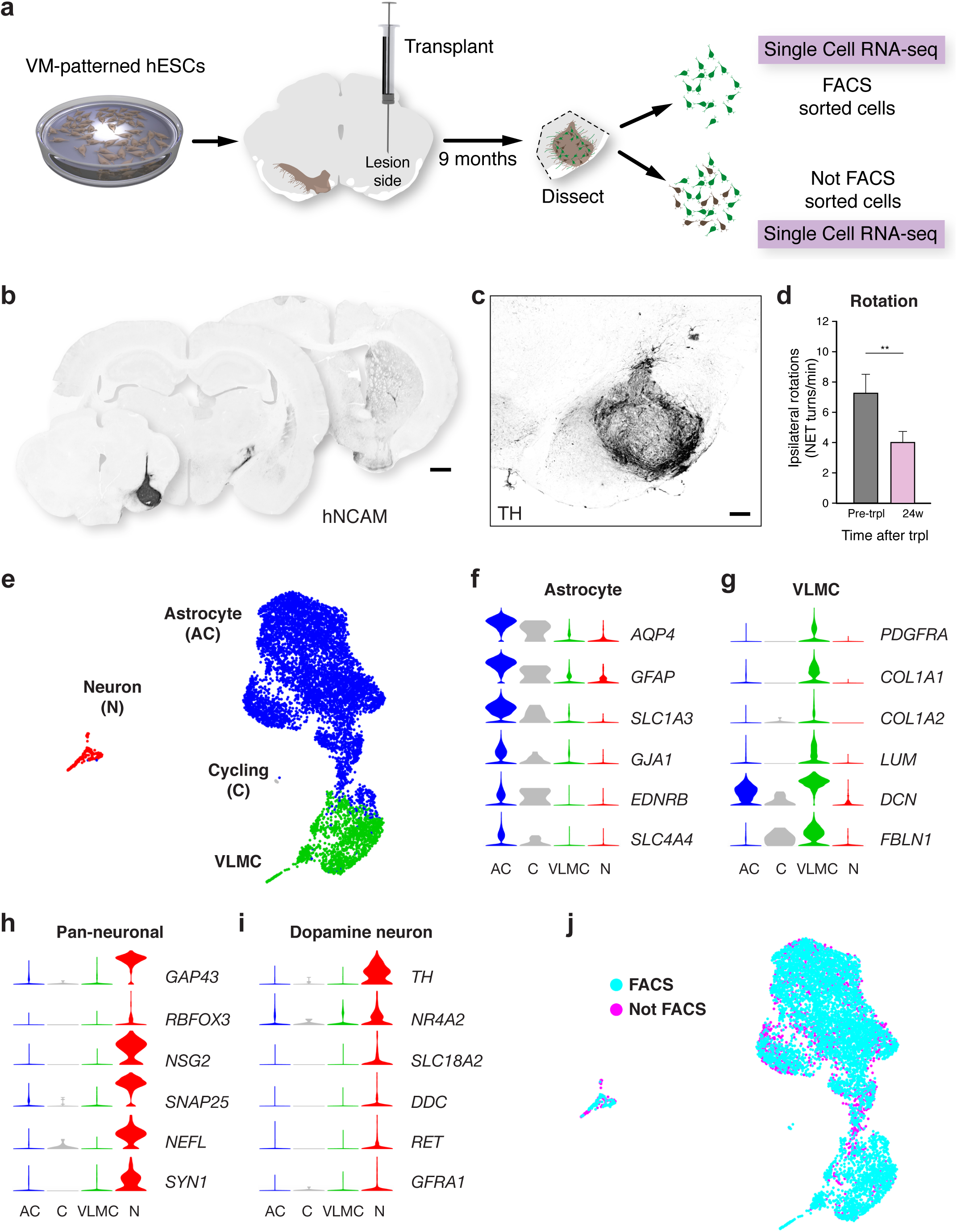
scRNA-seq analysis and histological validation of grafted cells into the midbrain. **a**, Schematic overview of experimental design. VM-patterned hESCs grafted to midbrain of 6-OHDA rats and analysed at nine months (n=6). These rats were used as follows: scRNA-seq n=3; histology n=3, functional recovery n=6. **b**, Overview of hNCAM fiber outgrowth from an intranigral transplant of hESC-DA neurons showing extensive re-innervation of the host striatum. **c**, Immunohistochemistry showing TH staining in graft core of a hESC-derived intranigral graft at 9 months post-transplantation. **d**, Drug-induced rotation test showing functional recovery in rats that have been transplanted to the midbrain with hESC-derived cells (n= 6 rats; mean ± SEM; p < 0.05 compared to post-lesion; two-tailed paired t-test). **e**, t-SNE showing clustering of analyzed cells after grafting to the midbrain. **f-i**, Expression level per cluster for indicated genes. Indicated genes are established markers for astrocytes, VLMCs, neurons and DA neurons, respectively. All indicated markers were significantly enriched in clusters marked as “Astrocyte”, “VLMC” and “Neuron”, respectively. **j**, t-SNE of grafted cells as shown in Fig. 4e. Cells isolated by FACS (blue) or not by FACS (magenta) are indicated. Scale bars, 1 mm (b); 200 µM (c).

Next, data from all hESC-derived grafted cells (in total 8558 cells) was integrated (683 cells grafted to the striatum; Smartseq2 sequencing and 7875 cells grafted to the midbrain; 10X Genomics) by using the Seurat data integration strategy^26^. Cells grafted to the striatum (sorted and subjected to Smartseq2 sequencing) and to the midbrain (sorted and unsorted, subjected to 10X Genomics) were all distributed in the main clusters (Extended Data Figures 9 and 10; see also enriched markers in Extended Data Table 5). Thus, VM-patterned hESC-derived cells formed the three major cell types (astrocytes, neurons and VLMCs) regardless of how they were isolated (direct isolation or FACS) and independent of grafting site (striatum or midbrain). In addition, similar to when cells were grafted to the striatum, the midbrain grafts resulted in a low yield of recovered neurons, but in the integrated data set the total number of analyzed neurons was substantially increased (from 35 to 232 cells) and the neuron cluster expressed *TH* and several other DA neuron markers including *NR4A2, SNCA, PBX1*, and *DLK1* (Extended Data Figure 10g). These markers were broadly distributed within the cluster and separate computational analysis of neurons alone did not give any indication of separate sub-clustering or marker expression for other neuron types, including serotonergic neurons (not shown). It thus seems likely that DA neurons (and DA neuron-like cells) are the dominating neuronal feature within grafts. However, additional analysis of larger number of recovered neurons will be important to further resolve graft neuron diversity in future studies.

The presented data provides a first unbiased assessment of graft composition from transplants of fetal VM and hESC-derived VM progenitors, which has been an outstanding question in the PD transplantation field for more than 30 years. Efforts to use stem cell-derived DA neurons instead of fetal VM tissue in clinical applications have progressed immensely and is now at the verge of entering clinical trials^3^. The results presented here have significance for transplanting proliferating hPSC-derived VM progenitors in patients in several different ways: *First*, scRNA-seq of cells after grafting demonstrated that only very few hESC-derived cells belonging to the astrocyte cluster remained proliferative. *Second*, serotonergic neurons that potentially can cause dyskinesias were not detected in the hESC-derived grafts (Freed et al. 2001 and Olanow et al 2003). Thus, the results presented here support the conclusion that these cells are safe to use in cell therapy. *Third*, scRNA-seq of fetal and hESC-derived grafted cells demonstrated that oligodendrocytes are only generated from fetal cells. Although the number of sequenced fetal cells was low, this finding was firmly validated in comprehensive histological analyses of several fetal and hESC-derived grafts. *Fourth*, the transcriptional profiling also revealed that hESC-derived grafts contained a cell type, that was previously not known to be part of the graft. These cells resembled normal vascular fibroblasts/VLMCs that have recently been identified in the mouse brain using scRNA-seq^15,21^. The graft-derived VLMCs were localized around blood vessels intermingled with endogenous vascular cells. The presence of graft-derived VLMCs could potentially be beneficial for graft survival, maturation and function by promoting a rapid vascularization and re-establishment of the blood brain barrier. They might also affect the immunological profile of the graft, and thus the immune response of the host, which is an important aspect to consider when designing and interpreting the first clinical trials using hESC-derived cells.

## Methods

### Animals

Athymic nude female rats (180g) were purchased from Harlan laboratories (Hsd:RH-Foxn1_rnu_) and housed in individual ventilated cages under a 12:12 hour dark-light cycle with *ad libitum* access to sterilized food and water. All procedures on research animals were in accordance with the European Union directive (2010/63/EU) and approved by the local ethical committee at Lund University as well as the Swedish Department for Agriculture (Jordbruksverket). For all surgical procedures rats (minimum 225 g/16-18 weeks) were anaesthetized via i.p. injections using a 20:1 mixture of fentanyl-dormitor (Apoteksbolaget). The animals were rendered hemiparkinsonian via unilateral injection of 4µL of the neurotoxin 6-hydroydopamine at a concentration of 3.5µg/µL (calculated from free-base HCl) aimed at the following stereotaxic coordinates (in mm): Anterior: −4.4, Medial: −1.2 and Dorsal: −7.8 with the incisor bar set at 2.4. Injections were performed as described^27^ at an injection speed of 1µL/minute with an additional 3 minutes allowed for diffusion before careful retraction of the needle. Drug-induced rotations were assessed four weeks post-lesion (d-amphetamine, 2.5mg/kg, i.p., Apoteksbolaget, Sweden) for 90 minutes in automated rotometer bowls (Omnitech Electronic Inc., USA). Rotational data was expressed as net turns per minute. The cylinder test was also used by placing animals in a glass cylinder (diameter 30cm) for 5 minutes while video recording. We counted the total number of touches the animals performed over the 300 second interval with the paw ipsilateral and contralateral to the lesion/graft, respectively. Paw use preference was expressed as bias score. After matching rats into groups according to their behavioural performance each rat received a single cell suspension of the respective hESC or human VM tissue. For intrastriatal grafts of hESCs-derived VM-patterend cells, we injected a total of 300.000 cells in 4 µL over four 1µL deposits (75.000 cells per deposit) at the following coordinates relative to bregma (in mm): Anterior: +1.2, Medial: −2.6, Dorsal: −4.5 and −4.0 and Anterior: +0.5, Medial: −3.0. For intranigral grafts of hESC-derived VM progenitors, we injected a total of 150’000 cells over 2 deposits (75.000 cells per deposit) to the midbrain at the following coordinates AP:-5.2; ML-2.3; DV-6.0/-6.5; TB flat head. At each deposit the injection-cannula was left in place for an additional 2 minutes before careful retraction to allow for settling of the tissue. The maturation of the graft was monitored using d-amphetamine induced rotations as described above. On the day before perfusion a second cylinder test was performed. After behavioral data had been collated, the rats were transcardially perfused using 50 ml physiological saline solution (8.9% Saline) over 1 minute followed by 250ml 4% PFA (pH7.4) solution over 5 minutes. For hESC-derived DA progenitors eight animals were transplanted in the intrastriatal grafting experiments. One rat had no graft and the remaining seven were used for analysis (two for sequencing and five for histology). For the intranigral transplants, nine rats were transplanted: one died after surgery, two were used to ensure good graft survival and maturation after six months and the remaining six were kept until the nine month endpoint. At this time three were used for sequencing and three for histology.

### Human Fetal Tissue preparation

Human fetal tissue from a legally terminated embryo was collected in accordance with existing guidelines with approval of the Swedish National Board of Health and Welfare and informed consent from women seeking elective abortions. To determine the gestational age of the embryo, the crown-to-rump length was measured and the embryo were staged according to week post-conception.

Fetal VM tissue was prepared for transplantation according to the same standard operating procedures (SoPs) as were approved for the clinical TRANSEURO trial (www.transeuro.org.uk). The fetal VM was dissected, cut into smaller pieces and incubated in TrypLE Select/Dornase alpha (DA, 20U/ml) for 20 min at 37°C for further dissociation. To achieve a crude cell suspension the tissue was then triturated first with a large tip (1000 µL) and then with a small tip (100 µL) in HBSS/Tirilazad Mesylate (TM)/DA. The cells where finally resuspended in 24 µL HBSS/TM/DA which is the same medium used for transplanting the hESC-derived progenitors. Each VM was transplanted to 3 rats. For the experiment used for sequencing, the 2 rats used to isolate grafts had reduced rotational scores (one from 6.29 to 0.14 and the other from 4.35 to −0.91) indicating functionality of the grafts. The dorsal midbrain was also dissected and was used to assess the viability of the fetal tissue.

### hESCs differentiation

hESCs (RC17: hPSCreg RCe021-A; H9: hPSCreg WAe009-A; HS980a: hPSCreg KIe033-A) and iPSCs (Miltenyi) were differentiated into VM-patterned progenitors using both our research-grade differentiation protocol (H9)^6,18^ and our GMP-grade protocol (RC17 and HS980a)^4^ and transplanted after 16 days of differentiation. Only cell preparations that could be approved according to a pre-determined quality range^4^were used for grafting. For expression of nuclear expression of GFP, RC17 hESCs (passage 25) were transduced with a lentiviral construct expression a histone H2B-GFP fusion protein under control of the human synapsin promoter (Addgene plasmid # 30456). The cells were transduced at a multiplicity of infection (MOI) of 6 while cultured on Laminin-521 in iPS-Brew (Miltenyi Biotec). For terminal differentiation, the VM-patterned progenitors were replated on day 16 as described^4^ and kept in culture until day 42-52 of differentiation.

To further assess the presence of VLMCs after *in vitro* differentiation we adopted two additional recently published/patented protocols that are intended for clinical use^11,25^ (see also https://patents.justia.com/patent/20180094242). For VM-patterned organoid differentiation RC17 hESCs were differentiated into self-organized organoids as described^28^, in presence of CHIR (CHIR99021) and SHH (SHH-C24II) for midbrain patterning^7^.

### Tissue dissociation and FACS sorting

Grafted human fetal ventral midbrain and grafted VM-patterned hESCs cells were dissected from rat striatum (n=2 for fetal VM intrastriatal grafts, n=2 for hESC intrastriatal grafts and 3 for intranigral grafts) and dissociated into a single cell suspension using the papain kit (Worthington) as previously described^29^. After dissociation, the cells were FACS sorted based on size (fetal tissue) or GFP expression (VM-patterned hESCs) using BD FACSAria III Cell Sorter (Extended Data Fig. 1a and b). The single cell suspension of human fetal ventral midbrain and VM-patterned hESCs before grafting were FACS sorted based on size. Tissue from individual animals were processed and sequenced separately.

### Library Preparation and Sequencing

The cDNA libraries from the sorted cells were generated using the Smartseq2 protocol^30^. cDNA libraries were tagmented as previously described, using a home-made Tn5 enzyme^31^ and Nextera dual indexes (i5 + i7). The quality of cDNA and tagmented cDNA was checked on a High-Sensitivity DNA chip (Agilent Bioanalyzer). Illumina HiSeq 2000 was used for sequencing, giving 43 base pair reads after de-multiplexing.

For 10X samples, single-cell suspensions were loaded onto 10X Genomics Single Cell 3′ Chips along with the reverse transcription (RT) mastermix per the manufacturer’s protocol for the Chromium Single Cell 3′ Library to generate single-cell gel beads in emulsion (GEMs). cDNA amplification was done as per the guidelines from 10X Genomics. Sequencing libraries were generated with unique sample indices (SI) for each sample with the following specifications Read1 28 cycles, Read2 98 cycles, Index1 8 cycles. Libraries for samples were multiplexed and sequenced on an Illumina NextSeq 500 machine using a 150-cycle NextSeq 500/550 High Output Reagent Kit v2. Sample Rat39_3a from Not FACS sorting was split into half (Rat39_3a and Rat39_3c) prior to cDNA amplification and was sequenced on separate lanes. Counts matrices obtained from these two samples were merged (rat39-3ac) during the downstream analysis.

### Read alignment and quality control

Reads for Smartseq2 data were aligned to the human genome (hg19) merged with *GFP* and TVA sequences using STAR v2.3.0^32^ and filtered for uniquely mapping reads. Gene expression was calculated as reads per kilobase gene model and million mappable reads (RPKMs) for each transcript in Ensembl release 69 using rpkmforgenes^33^. To exclude the contaminated rat cells, kallisto software was used for mapping of reads to the rat genome. The low-quality libraries were filtered out based on following parameters: < 5% uniquely mapping reads, > 25% fraction mismatches, < 10% exon mapping reads, >20% 3’mapping, < 3% or > 30% of all genes detected, < 10000 normalization reads, > 10% reads mapped to rat genome (Extended Data Fig. 1c). From 2206 sequenced cells 1406 passed the quality control.

For 10X Genomics samples, raw base calls were demultiplexed and converted to sample specific fastq files using cellranger mkfastq program. Raw reads were processed using cellranger count version 3 pipeline using default parameters. Precisely, this pipeline uses STAR to map cDNA reads to the transcriptome (GRCh38). Only exonic reads that uniquely mapped to transcriptome were used for the downstream analysis. Aligned reads were filtered for valid barcodes and UMI and observed cell barcodes were retained if they were 1-Hamming-distance away from entering into the whitelist of barcodes. Non-homopolymer UMIs with quality score greater than 10 were retained. Distribution of UMIs in each unique cell barcodes was examined and cell barcodes with UMI counts that fell within the 99^th^ percentile of the defined range of estimated cell count were selected. Cells were excluded if more than 10% reads aligned to Rat genome (Rnor6) (Extended Data Fig 8c). Low quality cells were filtered out based on: a) fraction of UMIs mapping to mitochondria larger than 0.15 and b) log10 detected genes per cell (nGene) below mean minus two standard deviations for each sample separately (Extended Data Fig 8b). A total of 7875 cells were used for downstream analysis.

### Smartseq2 cell population definition

Seurat package^34^ version 2.2.0 was used to normalize the data by gene detection and to extract the variable genes which were used for downstream analysis (x.low.cutoff = 1, x.high.cutoff = 10, y.cutoff =1). Only statistically significant principal components were included (Cells after grafting – PC: 1-12, Cells before grafting-PC:1:6, 8, 10:12). Clusters have been defined with Seurat function FindClusters using resolution 0.6. This gave five clusters for the cells before grafting (Extended Data Fig. 2a) and seven clusters for the cells after grafting (Extended Data Fig. 4a). After analysis of differentially expressed genes between the clusters, clusters with few differentially expressed genes between them and close proximity on the tSNE were manually defined as a single cluster, which gave 4 clusters for cells before grafting and 4 clusters for cells after grafting (Fig. 1e and 2a). The clustering results were visualized with the t-SNE plot (Rtsne: T-Distributed Stochastic Neighbor Embedding using a Barnes-Hut implementation, https://github.com/jkrijthe/Rtsne). To extract the cluster-specific genes we used the Samseq software package^12^. Differentially expressed genes for each cluster are in Extended Data table 1 and 2. Genes were considered significantly differentially expressed if q-value < 0.01.

### Integrated analysis

All clustering and analysis was performed with Seurat v3.0.0^26^. Cells from each rat graft was merged into individual datasets as well as grafted cells originating from hESCs in the SmartSeq2 dataset. Each dataset was log normalized and scaled with regression of number of features. The four datasets were integrated using the strategy described in Stuart et. al. using 30 principal components and 2000 anchor features. Dimensionality reduction was performed with UMAP (McInnes, L, Healy, J., UMAP: Uniform Manifold Approximation and Projection for Dimension Reduction, ArXiv e-prints 1802.03426, 2018) using 30 principal components. Cells were split into 4 clusters using Seurat default SNN clustering with resolution 0.1. Differential expression was performed with the Wilcoxon rank sum test (Extended Data Table 4 and 5). Genes were considered significantly differentially expressed if padj-value < 0.05.

### Pathway Analysis

The gene clusters after transplantation (genes from Extended Data table 2) were analyzed for biological functions using the Ingenuity Pathway Analysis program (QIAGEN Inc., https://www.qiagenbioinformatics.com/products/ingenuity-pathway-analysis). Only functional categories related to System Development functions were selected (Extended Data table 3) and ten most significant System Development functions of each cluster were plotted in Extended Data Fig. 6a.

### Comparison with other data sets

Gene expression profiles per cluster from the Mouse brain atlas^14^ was downloaded from their website (http://mousebrain.org). A marker gene set consisting of top 100 up-regulated gene per cluster among the cells after grafting (for either Smartseq2 data alone or integrated with 10x data), combined with marker genes for all the 256 cell types in the atlas was used in the comparison (Extended Data Fig. 6b, 6c, 11). The 256 atlas cell types were grouped into main clusters at Taxonomy rank 4 (39 groups) and mean expression per group was calculated using the marker gene set. These were compared to the mean expression in our clusters using spearman correlation.

### Cell Cycle Scores

G2M and S phase scores were calculated using the function CellCycleScoring in the Seurat package. As described^35^ the cells were classified as cycling if either of the scores were larger than 1. If scores were less than 1 they were defined as non-cycling.

### Immunohistochemistry

After perfusion, brains were post-fixed for 24 hours in 4% PFA and then cryopreserved in a 30% sucrose solution before being sectioned coronally on a freezing sledge microtome at a thickness 35µm in series of 1:8 or 1:12.

Immunohistochemistry was performed on free-floating sections and all washing steps were done with 0.1 M phosphate buffered saline with potassium (KPBS, pH = 7.4). The sections were washed three times and then incubated in Tris-EDTA pH8 for 30 min at 80°C for antigen retrieval. After washing an additional three times the sections were incubated in blocking solution (KPBS + 5 % serum of species the secondary AB was raised in + 0.25 % Triton X-100) for 1 hour, before adding the primary antibody solution. The primary antibodies used were: mouse anti-HuNu (1:200, Merck Millipore, cat. no. MAB1281), rabbit anti-TH (1:1000, Merck Millipore, cat. no. AB152), rabbit anti-NeuN (1:500, Merck Millipore, cat. no. ABN78), sheep anti-hCOL1A1 (1:200, R&D, cat.no. AF6220), rabbit anti-COL1A1 (1:200, Abcam, cat. no. ab34710), rabbit anti-hPDGFRa (1:300, Cell Signaling, cat. no. 5241), rabbit anti-Olig2 (1:500, Neuromics, cat. no. RA25081) and chicken anti-GFP (1:1000, Abcam, cat. no. ab13970). After incubation with primary antibodies over night at room temperature (RT), the sections were washed twice and incubated in blocking solution for 30-45 min. For fluorescent immunolabeling, the sections were then incubated with fluorophore-conjugated secondary antibodies (1:200, Jackson ImmunoResearch Laboratories) for 2 hours at RT, washed three times and then mounted on gelatin coated slides and cover slipped with PVA-DABCO containing DAPI (1:1000). For di-amino-benzidine (DAB) stainings, the sections were incubated with secondary biotinylated-horse antibodies (1:200, Vector Laboratories) for 1 hour at RT, washed three times and then incubated with avidin-biotin complex (ABC) for 1 hour at RT for amplification. Peroxidase driven precipitation of DAB was used to detect the primary antibody. In this step, sections were incubated in 0.05% DAB for 1-2 minutes before addition of 0.01% H_2_O_2_ for 1-2 minutes. After development of the DAB staining, the sections were mounted on gelatin coated slides and then dehydrated in an ascending series of alcohols, cleared in xylene and coverslipped with DPX mountant.

The hESC graft in Fig. 1b, 3b-f and Extended Fig. 1d-f has previously been analyzed for a different purpose and published^10,4,24^. The micrographs presented here are derived uniquely for this study.

### Immunocytochemistry

Terminally differentiated cell cultures (day 42-52) were fixed in 4% PFA for 15 min and then washed three times with PBS. For immunocytochemistry, the cells were blocked for 1-3 hours in PBS+5 % donkey serum + 0.1 % Triton X-100 before adding the primary antibodies solution. The primary antibodies included: rabbit anti-TH (1:1000, Merck Millipore, cat. no. AB152), sheep anti-hCOL1A1 (1:200, R&D, cat.no. AF6220) and rabbit anti-hPDGFRa (1:300, Cell Signaling, cat. no. 5241). After incubation with primary antibodies over night at 4°C, the cells were washed three times before adding fluorophore-conjugated secondary antibodies (1:200, Jackson ImmunoResearch Laboratories) and DAPI (1:500). The cultures were incubated with secondary antibodies for 2 hours and finally washed three times.

### Microscopy

Images were captured using either a flatbed scanner Epson Perfection V850 PRO, a Leica DMI6000B widefield microscope or a Leica TCS SP8 laser scanning confocal microscope. The image acquisition software was Leica LAS X and images were processed using Volocity 6.5.1 (Quorum Technologies) and Adobe Photoshop. Any adjustments were applied equally across the entire image, and without the loss of any information. The following images were digitally stitched from multiple images: Fig. 1a-b, Fig. 2h, Fig. 3a-f, Fig. 4, Extended Data Fig. 1d, Extended Data Fig, 5d, Extended Data Fig. 7a, c, m.

## Data availability

GSE118412 and GSE132758

## Acknowledgements

We thank Ulla Jarl, Marie Persson Vejgården and Mikael Sparrenius for technical assistance, Bengt Mattsson for his valuable help with microscopy and figure preparation and Prof. Anders Björklund and Dr. Laura Lahti for helpful comments and discussions. The research leading to these results has received funding from the New York Stem Cell Foundation (MP), the European Research Council (ERC Grant Agreement no. 30971, MP), the Swedish Research Council [grant agreements 521-2012-5624 (MP), 2016-02506 (TP) and 70862601/ Bagadilico)], Swedish Parkinson Foundation (Parkinsonfonden, MP), Swedish Brain Foundation (Hjärnfonden, MP), the Strategic Research Area at Lund University Multipark (Multidisciplinary research in Parkinson’s disease), Knut och Alice Wallenberg Foundation (TP), Söderbergs Stiftelse (TP) and Svenska Sällskapet för Medicinsk Forskning (KT). M.P. is a New York Stem Cell Foundation—Robertson Investigator.

## Author contributions

K.T. conceived the project, designed and performed scRNA-seq experiments, performed and interpreted computational analysis, analyzed all results in the project, and wrote part of the paper; S.N. conceived the project, designed and performed histological and *in vitro* experiments, interpreted histological data, analyzed all results in the project, and wrote part of the paper; Å.K.B and YS. performed and interpreted computational analysis; A.F. designed and performed *in vivo* experiments, scRNA-seq experiments and designed and performed histological experiments and *in vitro* differentiation experiments, interpreted histological data and analyzed results; A.H designed and performed *in vivo* experiments; D.H., A.A. and TC performed *in vivo* experiments; M.B performed experiments with fetal tissue, L.G., H.L.-M. and N.V. performed wet lab procedures for scRNA-seq experiments; A.K. conceived the project, designed experiments and interpreted results. M.P. and T.P. conceived the project, designed experiments, interpreted results and wrote the paper with input from all authors.

