## Supplemental figures 1-11 for "Single Cell Gene Expression Analysis Reveals Human Stem Cell-Derived Graft Composition in a Cell Therapy Model of Parkinson’s Disease"

#### Extended data figure 1

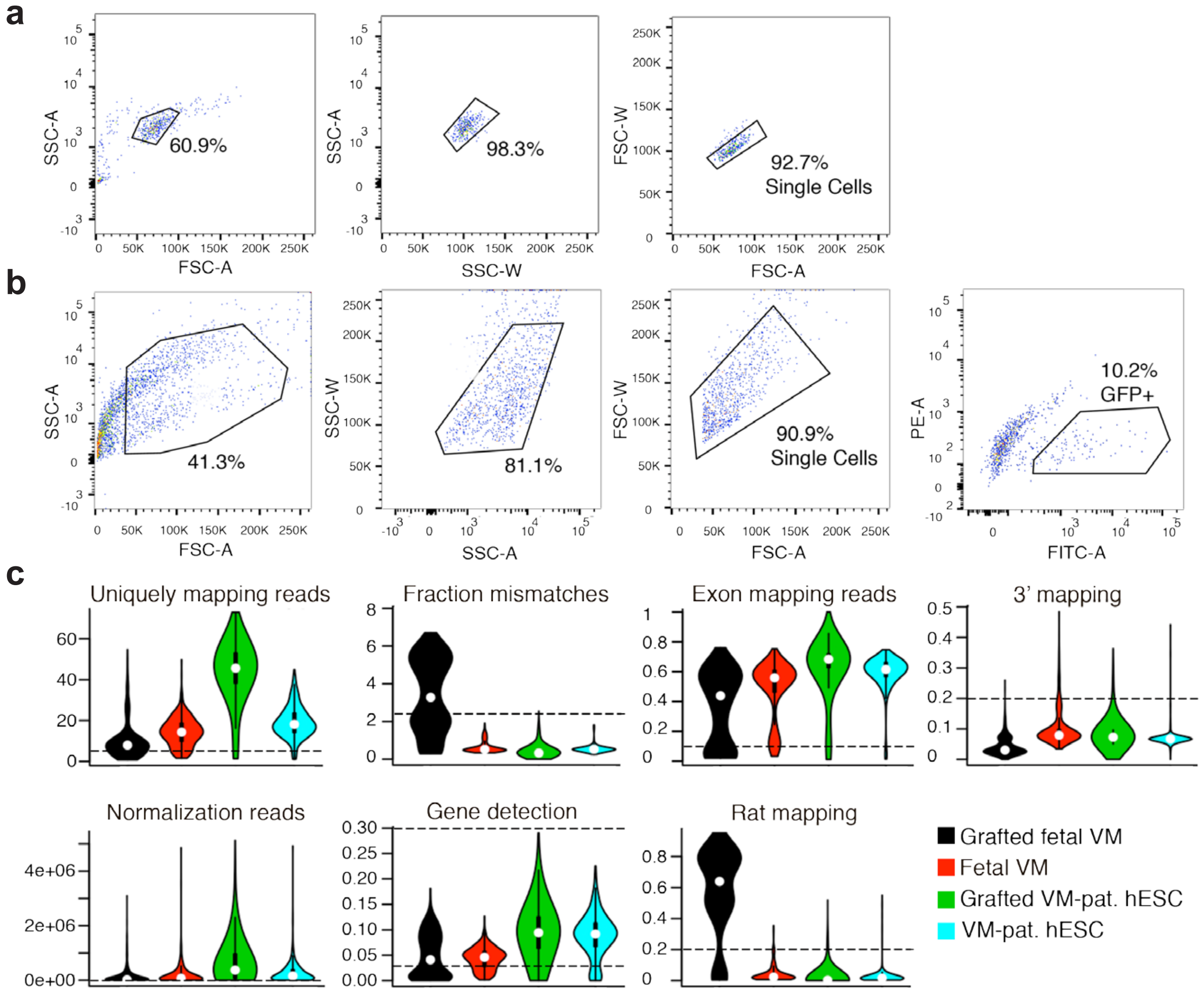


**d**

| | Fetal VM | VM-pat. hESC | Grafted fetal VM | Grafted VM-pat hESC |
|---|---|---|---|---|
| Single cell RNA-seq | 384 | 480 | 480 | 862 |
| Cells after QC | 256 | 404 | 63 | 683 |

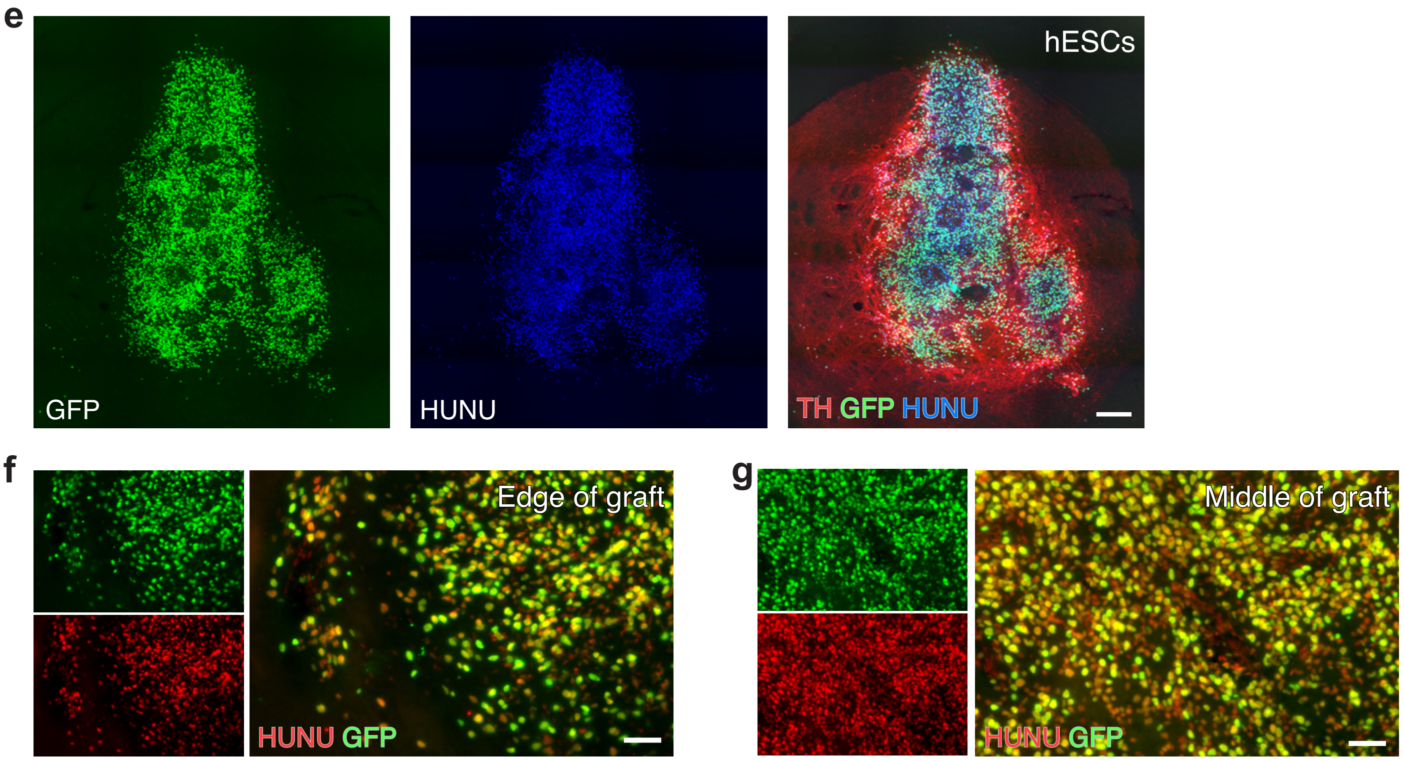

#### Extended data figure 2

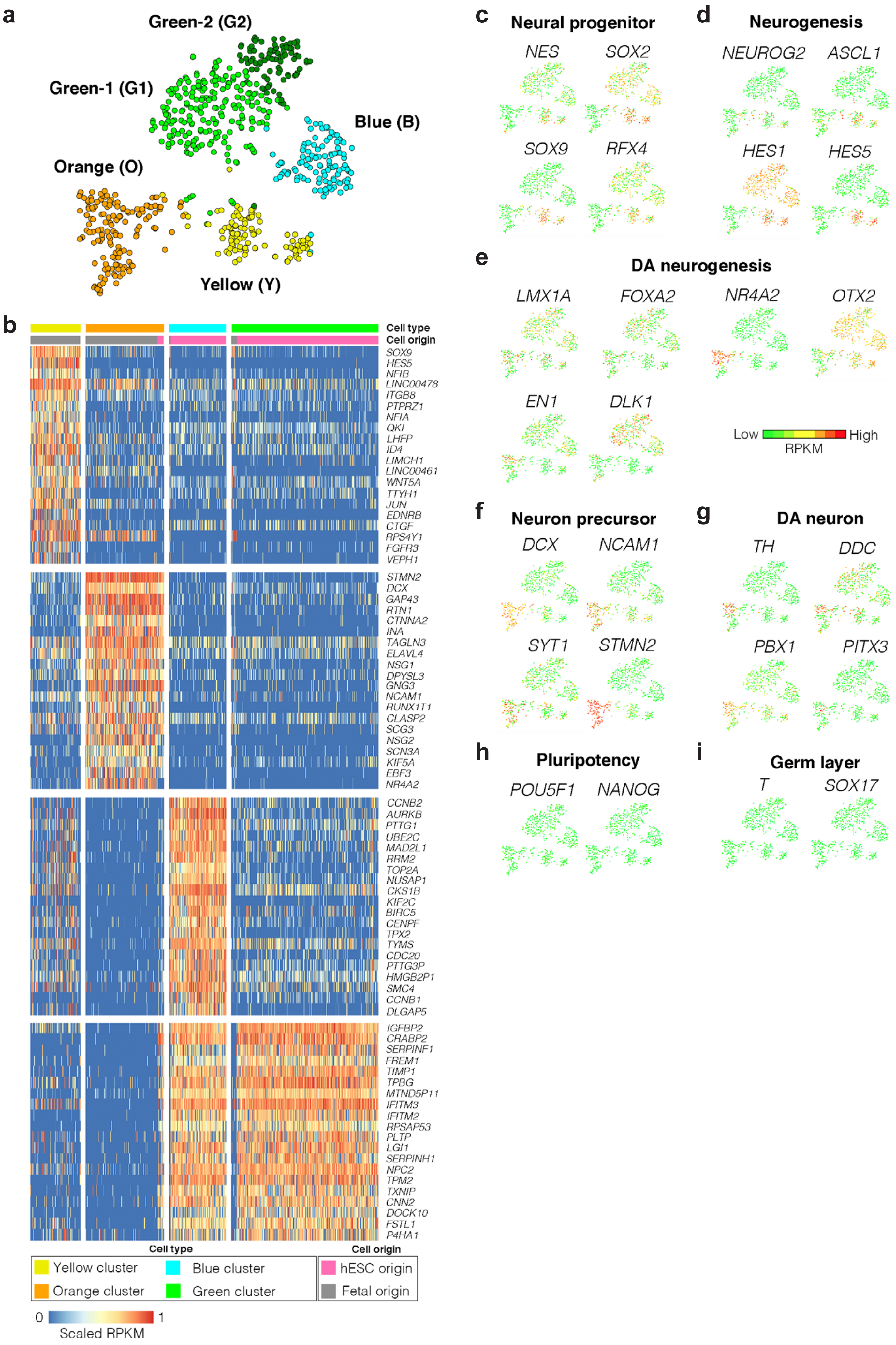

#### Extended data figure 3

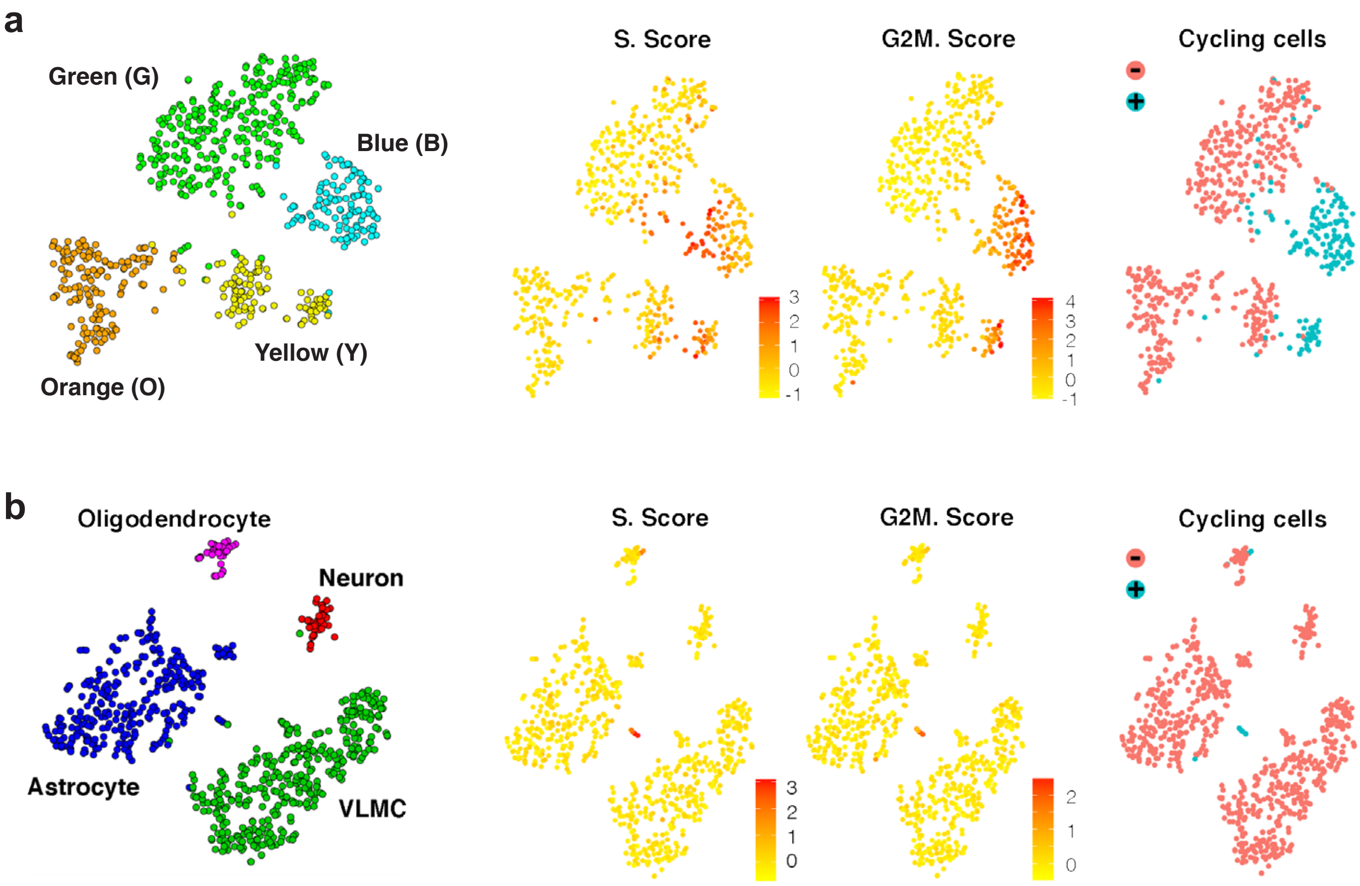

#### Extended data figure 4

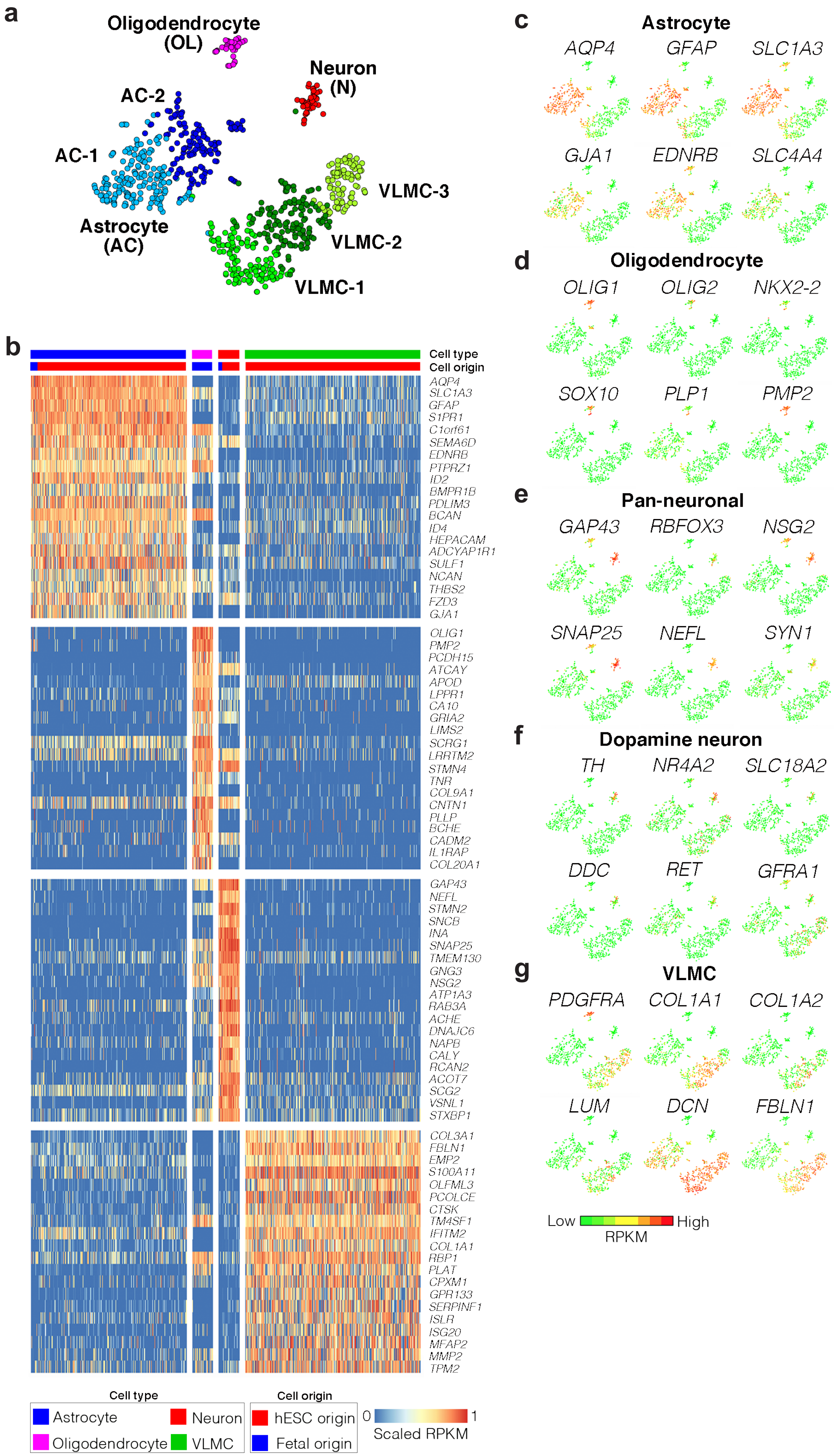

**Extended data figure 5**

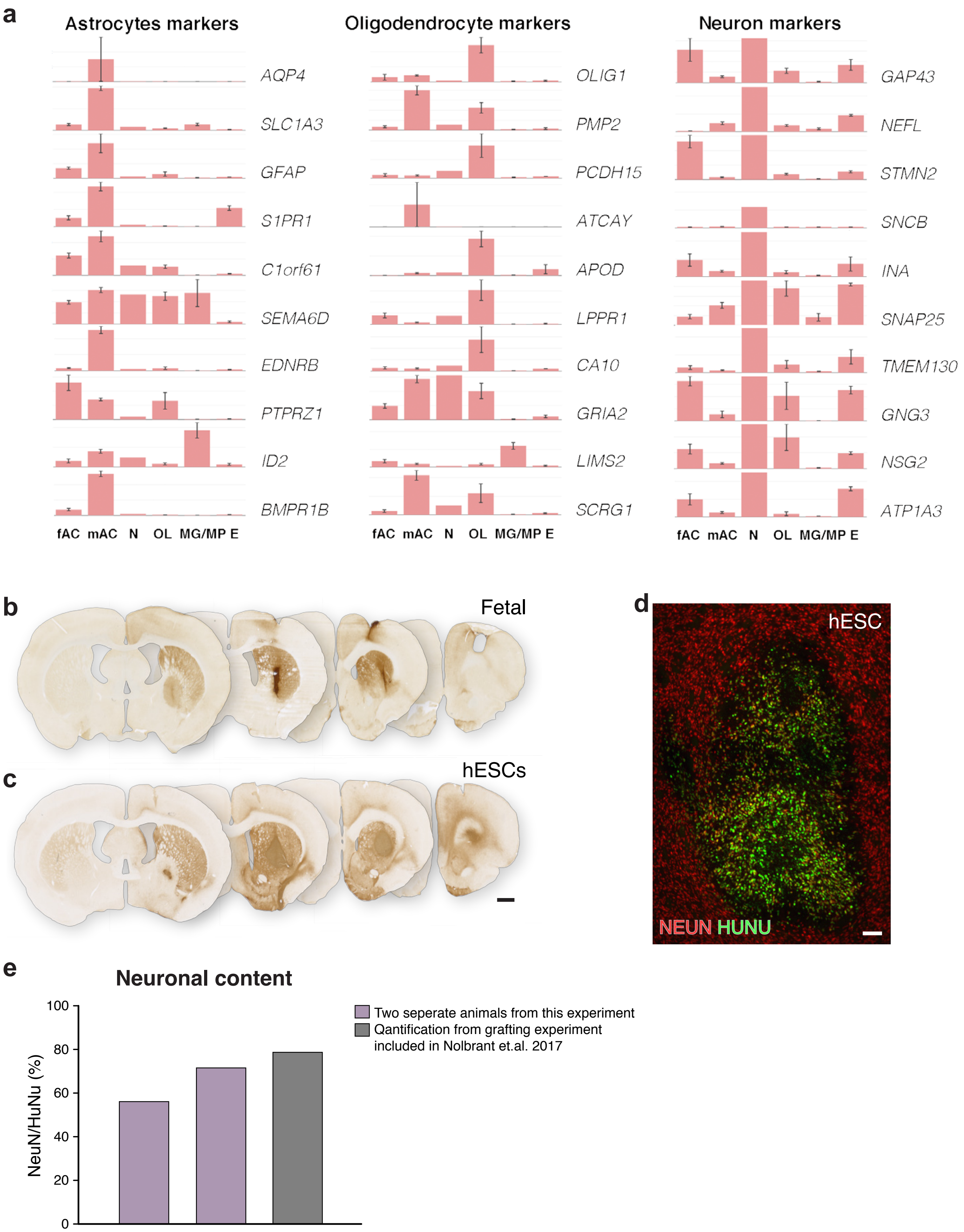

#### Extended data figure 6

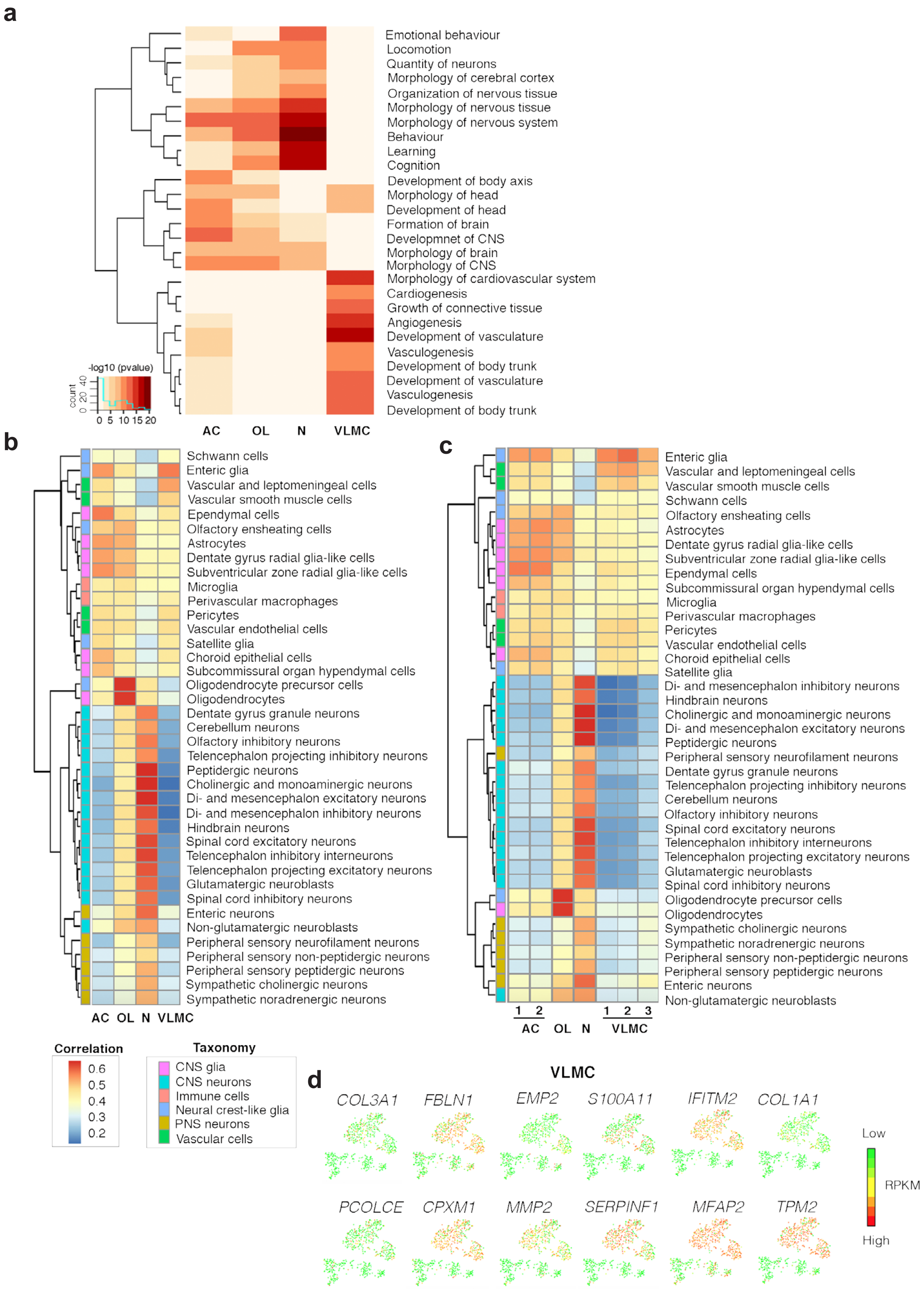

Extended data figure 7

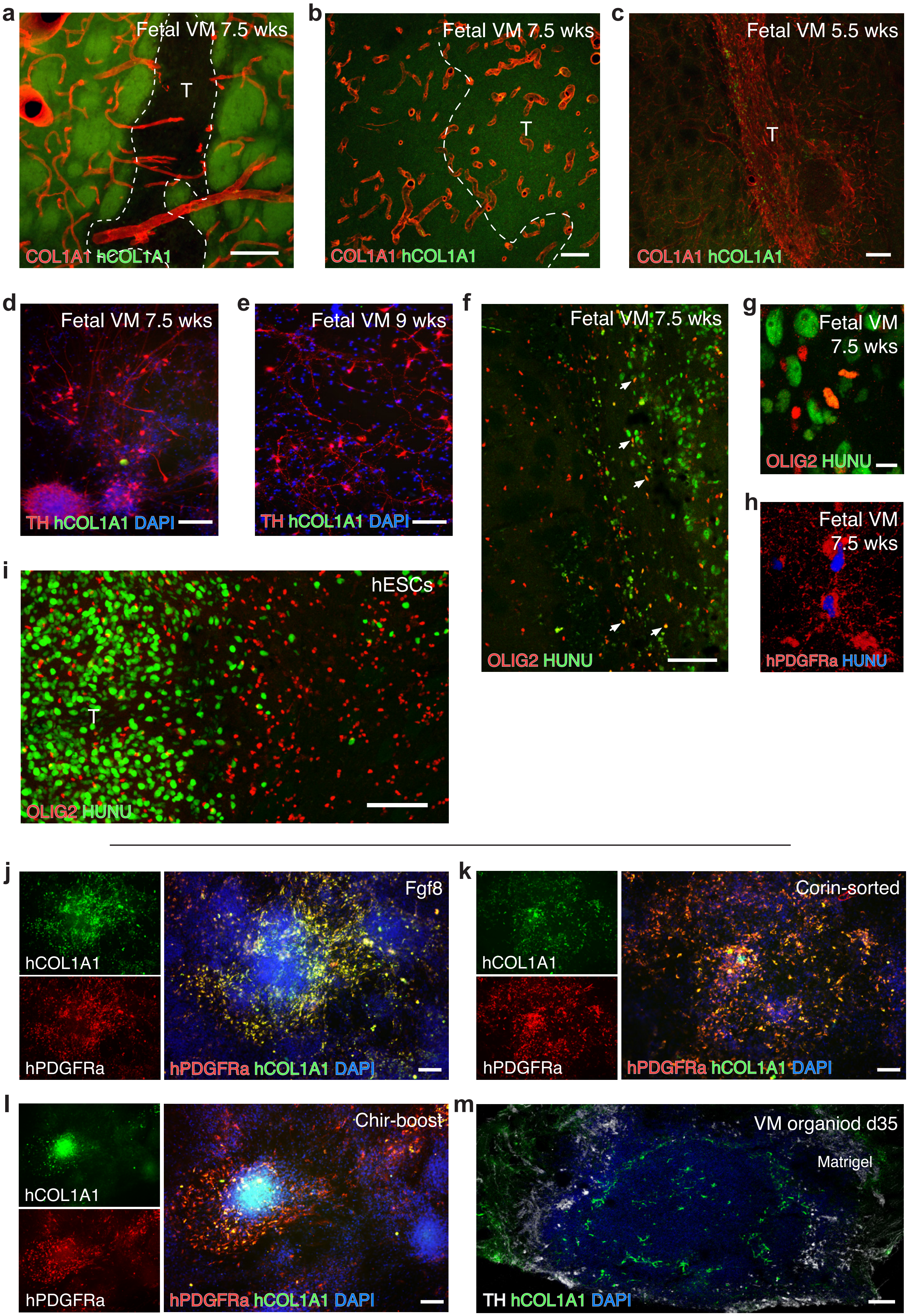

#### Extended data figure 8

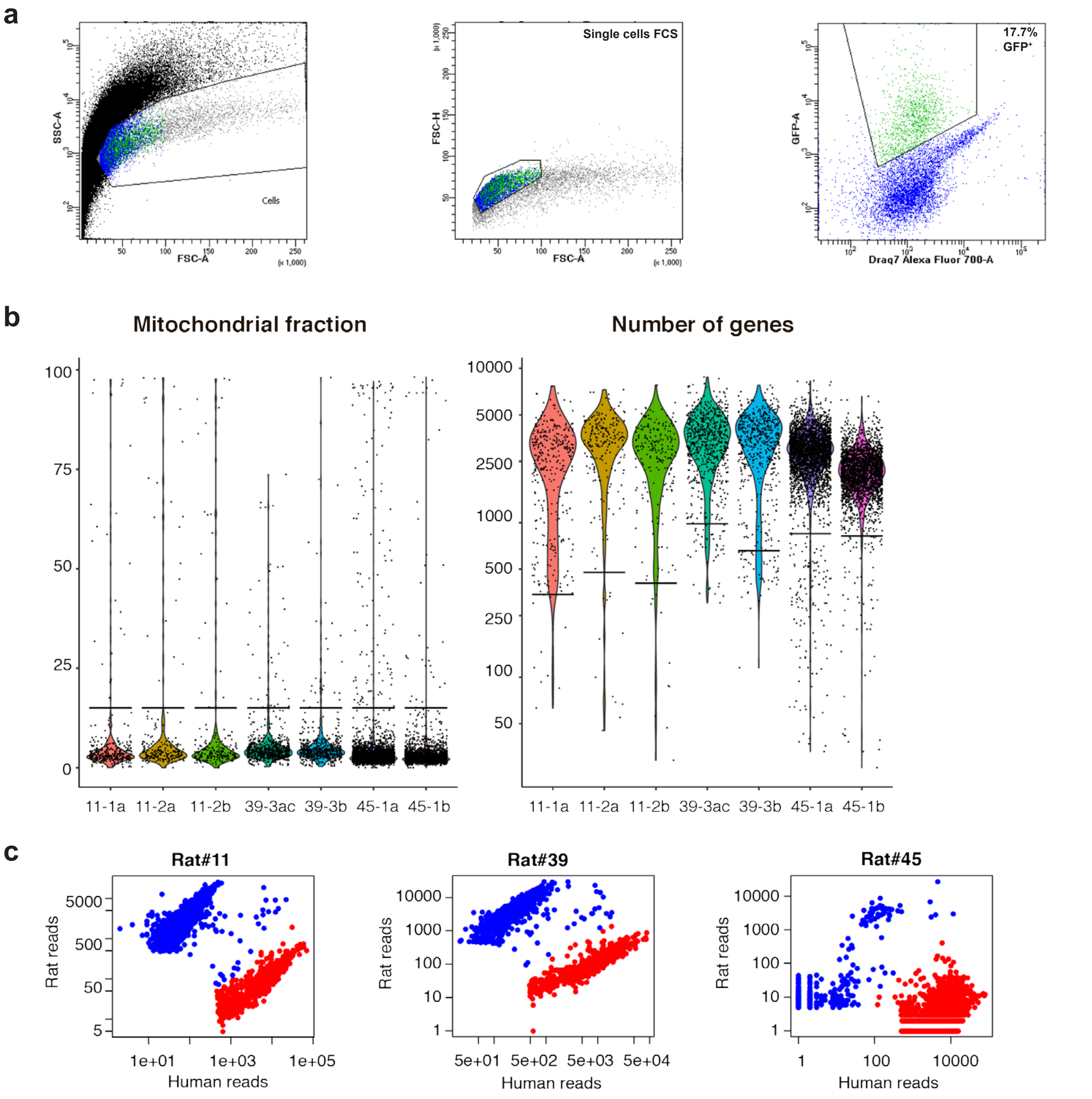

**Extended data figure 9**

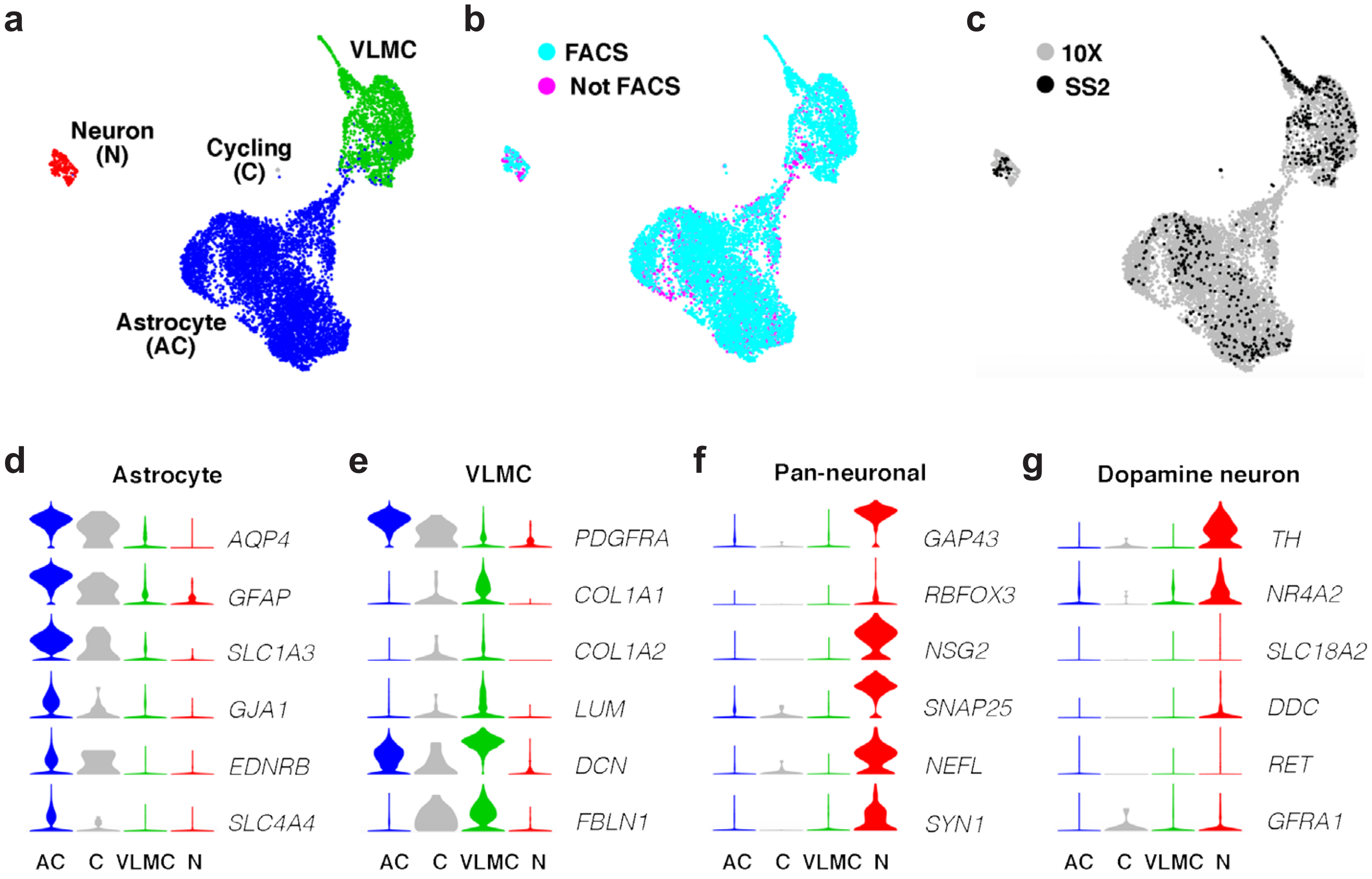

Extended data figure 10

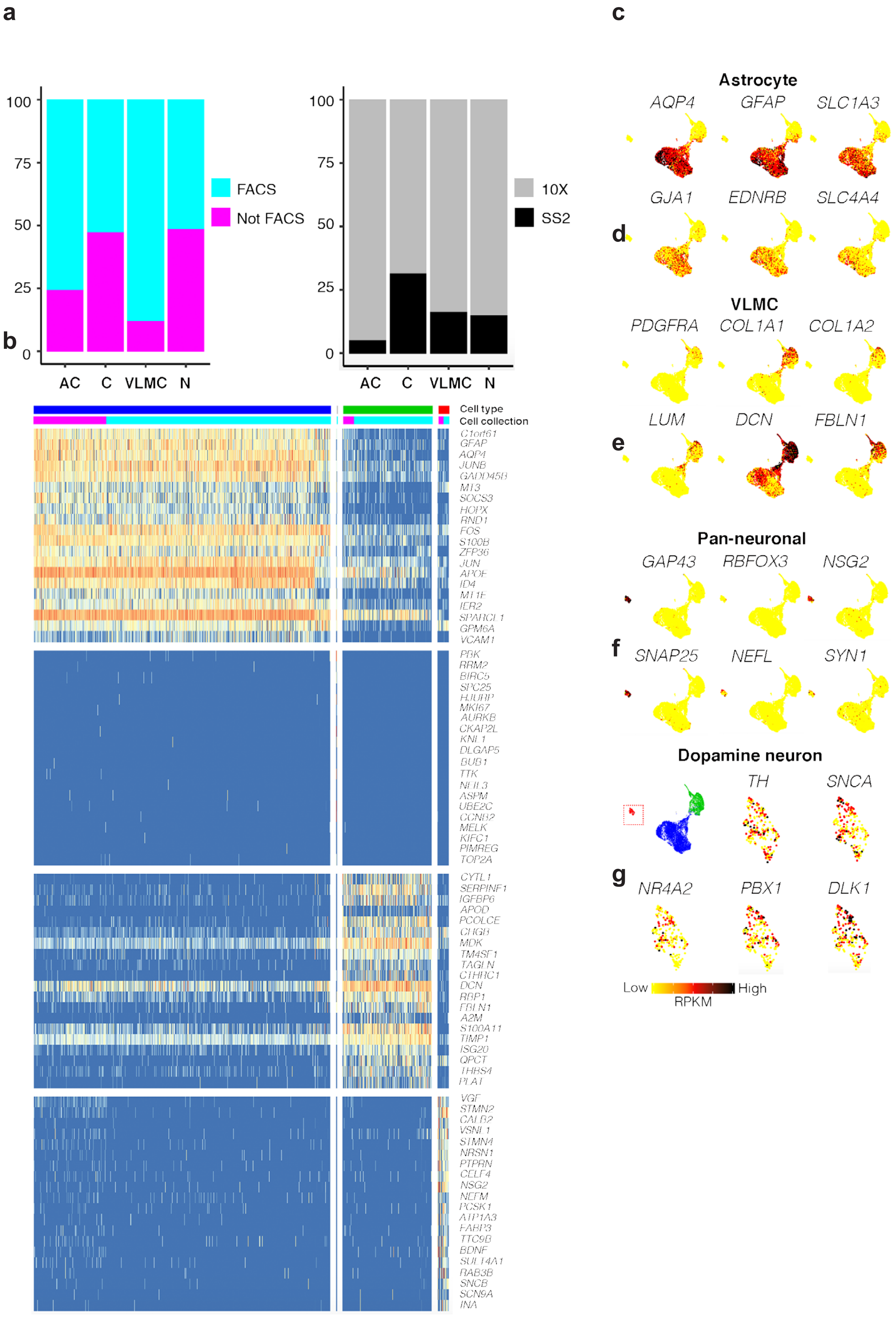

### Extended data figure 11

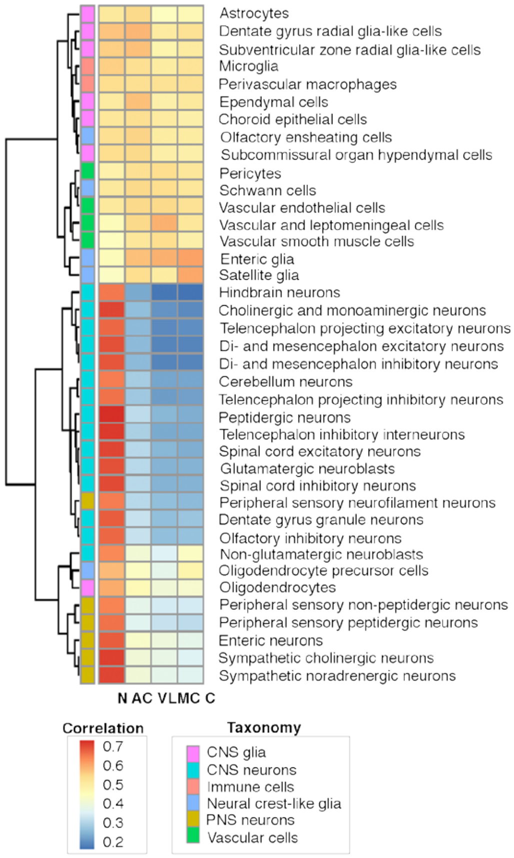

**Extended Data Figure 1**

**Isolation of cells by FACS and histological validation of GFP expression**

**a,** FACS strategy for sorting of the single cells based on the cell size. FSC-A: Forwards Scatter area; FSC-W: Forward Scatter width; SSC-A: Side Scatter area; SSC-W: Side scatter width; FITC-A: Area of fluorescent dye fluorescein; PE-A: Area of fluorescent dye phycoerythrin.

**b,** FACS strategy for sorting of the single cells based on the GFP expression. Abbreviations as in a.

**c,** Filtering of low quality cells based on specified parameters.

**d,** Table indicating the number of single cells used for scRNA-seq and single cells which passed the QC and were included for further analysis.

**e-g,** Immunohistochemistry showing GFP (from the reporter) and HUNU staining in the grafts 6 months after transplantation, at the graft edge (e) and in the graft core (f).

Scale bars, 250 μM (e) and 50 μM (f and g).

**Extended Data Figure 2**

**Analysis of scRNA-seq data from cells before grafting**

**a,** t-SNE showing clustering of all analyzed cells before grafting. Five clusters were defined with Seurat FindClusters. After analysis of differentially expressed genes and visualization by t-SNE, cells were manually defined as 4 clusters as indicated in Figure 1e and described in Methods.

**b,** Heatmap visualizing expression of top-enriched genes in clusters of the cells before grafting.

**c-i,** Expression of markers visualized on the t-SNE plot. The colors indicate the RPKM values.

**Extended Data Figure 3**

**Cell cycle scores in cells before and after grafting**

**a,** t-SNE plot showing clusters of cell types before grafting, cell cycle scores of the analyzed cells (S.Score and G2M.Score) and cycling cells (blue color). G2M and S cell cycle scores were calculated using the function CellCycleScoring in the Seurat package.

**b,** t-SNE plot showing clusters of cell types after grafting, cell cycle scores of the analyzed cells (S.Score and G2M.Score) and cycling cells (blue color).

**Extended Data Figure 4**

**Analysis of scRNA-seq data from grafted cells into the striatum**

**a,** t-SNE showing clustering of all analyzed cells after grafting. Seven clusters were defined with Seurat FindClusters. After analysis of differentially expressed genes and visualization by t-SNE, cells were manually defined as 4 clusters as indicated in Figure 2a and described in Methods. The clusters were assigned to different cell types as described for Figure 2.

**b,** Heatmap visualizing expression of top-enriched genes in clusters of the grafted cells.

**c-g,** Expression of markers visualized on the t-SNE plot. The colors indicate the RPKM values.

**Extended Data Figure 5**

**Validating cluster assignments and quantification of neurons**

**a,** Analysis of expression of top-enriched genes from clusters of the grafted cells in the RNA-seq transcriptome and splicing database (www.brainrnaseq.org). fAC – fetal astrocytes, mAC – mature astrocytes, N – neurons, OL – oligodendrocytes, MG– microglia, MP – macrophage, E – endothelial.

**b-c,** Overview of the pattern of hNCAM$^+$ axonal outgrowth derived from fetal- (b) and hESC-derived (c) intrastriatal grafts 6 months post-transplantation.

**d**, Representative HUNU/NEUN double immunostaining in hESC-derived grafted cells.

**e,** Estimation of the NEUN$^+$ fraction of HUNU$^+$ graft cells from 2 animals in this experiment and 1 animal from a previous experiment (Nolbrant, et al., (2017). *Nature Protocols*, *12*(9), 1962–1979). All cells in two sections (graft core and graft edge) per animal were quantified.

Scale bars, 1 mm (b and c) and 100 µM (d).

**Extended Data Figure 6**

**Analysis of VLMC functional pathways and comparison with brain clusters**

**a,** Top-significant Physiological System Development and Function categories from clusters of the grafted cells (Fig. 2a) as analyzed by Ingenuity Pathway Analysis (QIAGEN Inc., https://www.qiagenbioinformatics.com/products/ingenuity-pathway- analysis).

**b-c,** Correlation of clusters after grafting (horizontally) and mouse brain atlas (www.mousebrain.org) (vertically) as described in Methods.

**d,** Expression of VLMC markers visualized on the t-SNE plot of cells before grafting. The colors indicate the RPKM values.

**Extended Data Figure 7**

**Validation of VLMCs in hESC-derived grafts and in cultured VM-patterned hESCs**

**a-c,** Immunohistochemical analysis of COL1A1 and hCOL1A1 in fetal-derived grafts originating from 7.5 week old (a-b) or 5.5 week old VM tissue (c). Graft core is delineated with dotted line.

**d-e,** Fetal cultures dissected from 7.5 week (d) or 9 week (e) old VM tissue double immunostained for TH/ hCOL1A1.

**f-h,** Fetal grafts double immunostained for HUNU/OLIG2 (f-g) and hPDGFRa/HUNU (h).

**i,** hESC-derived grafts double immunostained for $OLIG2^{+}$/$HUNU^{+}$.

**j-l,** Representative pictures of PDGFRa /hCOL1A1 double immunostaining in terminally differentiated hESC *in vitro* cultures derived by three different clinically relevant VM-patterning differentiation protocols: the protocol used in this study (j), a protocol developed in the Studer lab that uses CHIR boost instead of FGF8 for proper caudalization (https://patents.justia.com/patent/20180094242)(k), and a protocol developed by the Takahashi lab where the cells are sorted based on CORIN prior to grafting (Doi, D., et al., (2014) *Stem Cell Reports*, *2*(3), 337–350; Kikuchi, et al., (2017) *Nature Publishing Group*, *548*(7669), 592–596). Nuclei were counterstained with DAPI.

**m,** TH/hCOL1A1 double immunostaining in self-organized midbrain patterned organoids.

Scale bars, 100 µM (a-f, i-m), 10 µM (g-h). T=transplant

**Extended Data Figure 8**

**Isolation of midbrain grafted cells by FACS and quality control of sequenced cells**

**a,** FACS strategy for sorting of the single cells based on GFP expression.

**b,** Filtering of low quality cells based on mitochondrial fraction and number of detected genes per cell.

**c,** Filtering of cells based on alignment to the rat genome.

**Extended Data Figure 9**

**Integrated data analysis of grafted cells into the striatum and midbrain**

**a-c,** t-SNE showing clustering (a), sorting (b) and type of library preparation and sequencing (c) of cells grafted into the striatum and midbrain.

**d-g,** Expression level per cluster for indicated genes. All indicated genes are enriched and known markers for astrocytes, VLMCs pan-neuronal cells and dopaminergic neurons.

**Extended Data Figure 10**

**Analysis of scRNA-seq data from grafted cells into the striatum and midbrain**

**a,** Graphs representing proportion of cell types per sorting and type of library preparation and sequencing.

**b,** Heatmap visualizing expression of top-enriched genes in clusters of the grafted cells.

**c-d,** Expression of markers visualized on the t-SNE plot. The colors indicate the RPKM values.

**e,** Expression of markers in the neuron cluster (cells marked in red) visualized on the t-SNE plot. The colors indicate the RPKM values.

**Extended Data Figure 11**

**Comparison of clusters after grafting into the striatum and midbrain with brain clusters**

Correlation of clusters after grafting into striatum and midbrain (horizontally) and mouse brain atlas (www.mousebrain.org) (vertically) as described in Methods.
